# Integration of gene expression with computational modeling reveals region-specific brain energy metabolism dysregulation in schizophrenia

**DOI:** 10.64898/2025.12.22.695881

**Authors:** Ilona Mäkinen, Marja-Leena Linne, Ole A. Andreassen, Tuomo Mäki-Marttunen

## Abstract

Schizophrenia is a highly heritable psychiatric disorder with suggested disturbances in multiple neurobiological pathways including brain energy metabolism, which is essential for sustaining precise neuronal signaling. The relationship between gene expression alterations and energy metabolism defects, however, is poorly understood. Here we combined differential expression (DE) analysis with computational modeling to investigate how the expression of cytosolic energy metabolism-associated genes influences the concentrations of key energy metabolites. The DE analysis used human post-mortem RNA sequencing data from the anterior cingulate cortex (ACC) and dorsolateral prefrontal cortex (DLPFC) from the CommonMind Consortium. Metabolic simulations were carried out using a biophysical model of brain energy metabolism (Winter et al. 2018). We conducted both population-average and subject-specific simulations and assessed the effects of DE genes individually and collectively.

The DE analysis identified nine genes in the ACC and 24 in the DLPFC. Approximately two thirds of the DE genes were downregulated in schizophrenia in both brain areas. Decreased expression of neuronal *LDHB* in the ACC caused a decrease of pyruvate and an increase of NAD^+^/NADH ratio. Furthermore, decreased expression of neuronal *PFKM* in the DLPFC caused an increase of glucose and NAD^+^/NADH ratio and a decrease of pyruvate, lactate, and ATP. These findings provide evidence for region-specific dysregulation of brain energy metabolism in schizophrenia and suggest mechanistic links between gene expression changes and altered levels of key energy metabolites.

**AUTHOR SUMMARY:** Schizophrenia is a mental disorder that affects perception, thought, and behavior, but its disease mechanisms are still not fully understood. One important aspect of brain function is energy metabolism, which powers nearly all cellular processes and supports communication between neurons. In this study, we explored how changes in the expression of specific genes involved in energy production influence the levels of key molecules in the brain. We combined human gene expression data from two brain regions with computer simulations of neuronal function. We found that certain genes had a strong impact on molecules such as glucose, pyruvate, lactate, ATP, and NAD^+^/NADH balance. Importantly, the effects were specific to particular brain regions. Still, the majority of gene expression changes had non-significant effects. By connecting gene expression with brain energy metabolism simulations, this work provides new insights into the biological mechanisms of schizophrenia and may guide future research aimed at understanding and improving brain function in the disorder.

## INTRODUCTION

Schizophrenia is a complex and heterogenous mental disorder with an estimated global lifetime prevalence of 0.4 % and lifetime morbid risk of 0.7 % [1]. The disorder is characterized by a variety of symptoms such as hallucinations, delusions, and disorganized speech (positive symptoms), and lack of pleasure and motivation, and social withdrawal (negative symptoms) [2,3]. Schizophrenia is also associated with cognitive decline including deficits in attention and memory [2,3]. Although the disorder has been extensively studied for decades, its etiology remains unclear. However, it is now widely regarded as a neurodevelopmental disorder influenced by environmental, social and genetic factors [3,4]. The heritability of schizophrenia is estimated to be around 80 % according to twin studies [5,6]. In addition, genome-wide association studies (GWAS) [7,8] have shown the polygenic nature of the disorder. GWAS have elucidated that schizophrenia is associated with a large number of common single nucleotide polymorphisms (SNPs) with small effect sizes, whereas other genomic factors like rare copy number variations (CNVs) with higher effect sizes have a much smaller contribution [9,10]. However, since GWAS associations are difficult to map to specific genes, transcriptomic analyses are essential for a broader understanding of schizophrenia’s disease mechanisms. Transcriptomic studies, alongside *in vitro*, animal, and imaging studies, suggest disturbances in several biological pathways of the brain. Furthermore, converging evidence suggests that dysregulation of brain energy metabolism and increased oxidative stress are key factors associated with schizophrenia [11–13].

The brain is one of the most metabolically active organs of the human body. It relies on a continuous turnover of adenosine triphosphate (ATP) to sustain appropriate neuronal functioning including maintenance and restoration of ion gradients, and uptake and recycling of neurotransmitters [14,15]. Dysregulation of brain bioenergetics has been associated with multiple brain diseases, and it can lead to impaired neurotransmitter synthesis and release, reduced synaptic function and plasticity, and accelerated neurodegeneration [16]. Disturbed brain energy metabolism has also been linked to schizophrenia, and especially to the negative and cognitive symptoms of the disorder [17,18]. Various findings such as insulin resistance, impaired glucose metabolism, elevated lactate levels, lower ATP levels, and disrupted redox balance, point to dysregulation of energy metabolism and oxidative stress in schizophrenia [13,17,19]. Since all current antipsychotics target mainly the positive symptoms of schizophrenia [20], gaining deeper knowledge on the mechanisms behind negative and cognitive symptoms is essential for developing more effective treatments. However, it has been challenging to decipher whether the bioenergetic disruptions of schizophrenia are caused by genetics or transcriptomics, or if they are a product of other pathophysiological processes (such as synaptic and neurotransmitter dysfunction), disease progression or antipsychotic use.

One promising method for studying the connection between transcriptomics and brain energy metabolism in disorders like schizophrenia is computational modeling [21]. It allows fast, cost-effective and reproducible way to study how gene expression alterations influence the function of energy metabolism processes and pathways. Modifying model parameters to reflect disease-specific gene expression profiles enables the investigation of their effects on transport dynamics, metabolic fluxes, and metabolite concentrations in response to neuronal activity. The first biophysically detailed model combining energy metabolism with neuronal activity and hemodynamics was developed by Aubert and colleagues in 2001 [22]. In the following years this model was refined by applying it to functional neuroimaging [23,24] and including the neuron-astrocyte compartmentalization and their functional interactions [25]. Building on the models developed by Aubert et al. [22–25], other researchers have refined brain energy metabolism models in various ways: for example, Cloutier et al. incorporated glycogen dynamics in astrocytes [26], while Winter et al. integrated the pentose phosphate pathway (PPP) into the model [27]. All the aforementioned models are compartmentalized brain energy metabolism models consisting of a series of coupled differential equations. The use of biophysically detailed models offers many advantages: it helps to understand and interpret *in vitro* and imaging data and allows detailed inspection of complex metabolic pathways. Lastly, computational modeling can be used for testing hypotheses and predicting outcomes on how transcriptional disturbances influence energy metabolism of the brain in health and disease.

In the present study we analyzed RNA sequencing (RNA-seq) data from anterior cingulate cortex (ACC) and dorsolateral prefrontal cortex (DLPFC), provided by CommonMind Consortium [28], to study the differential expression (DE) of cytosolic brain energy metabolism-related genes between schizophrenia patients (SCZ) and healthy controls (HC). Throughout this article, the abbreviation SCZ refers specifically to the schizophrenia subjects of the CommonMind data. We used a model of cytosolic brain energy metabolism developed by Winter et al. [27] to simulate the effects of altered gene expression on the concentrations of central energy metabolites: ATP, glucose, pyruvate, lactate and NAD^+^/NADH ratio. We analyzed how changes in gene expression affect both baseline metabolite levels and their time-dependent responses to neuronal activity. The simulations were performed using both population-average and subject-specific gene expression data, and both individual and collective effects of the DE genes were investigated. The results revealed that decreased expression of *LDHB* lead to decreased pyruvate levels and increased nicotinamide adenine dinucleotide (NAD^+^/NADH) ratio in the neurons in the ACC, and decreased expression of *PFKM* lead to increased glucose and NAD^+^/NADH and decreased lactate and pyruvate in both neurons and astrocytes, as well as decreased ATP in astrocytes in the DLPFC. By combining RNA-seq data with a biophysically detailed model of brain energy metabolism, this study provides a novel framework to link gene expression changes to metabolic alterations, offering new insights into the molecular mechanisms underlying schizophrenia.

## MATERIALS AND METHODS

### Differential expression analysis and imputation of cell type-specific gene expressions

A DE analysis was performed to study the expression alterations of cytosolic energy metabolism-related genes in schizophrenia. The data used in the DE analysis is human post-mortem bulk RNA-seq data from ACC and DLPFC, provided by CommonMind Consortium [28]. The ACC data consists of 251 HC and 230 SCZ, and the DLPFC data consists of 295 HC and 263 SCZ.

First, KEGG REACTION [29] and UniProt [30] databases were used to match the transport and metabolic reactions of the Winter model to their corresponding proteins and genes (S1 Appendix). We found 85 genes corresponding to the proteins of the model, and these genes were the subject of the DE analysis. The analysis was performed using R Statistical Software (v4.4.1; R Core Team 2021) [31]. Ensembl gene IDs used in the CommonMind data were matched to HUGO gene names using AnnotationDbi (v1.66.0) and org.Hs.eg.db (v3.19.1) packages. Before the DE analysis, the genes whose read count across all samples was in the lowest 10 % were filtered out. The DE analysis was performed using DESeq2 package (v1.44.0) [32] and its *DESeq* function which uses Wald test to calculate statistical significance. In addition to the pre-filtering, additional filtering of lowly expressed genes was performed by the *results* function of the DESeq2 package. The *results* function also accounts for multiple testing by using Benjamini-Hochberg correction. The significance threshold was set at α = 0.01. The design formula used in the DESeq2 analysis was

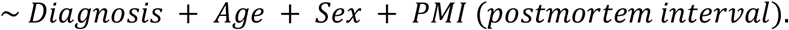

Some of the DE genes (DEGs) had to be left out from subsequent analyses because the Winter model does not describe their effects on energy metabolism or because there is not sufficient evidence on their protein-level expression in the brain. Only genes that could be modeled and had sufficient evidence of protein expression were included in the subsequent analyses (S2 Appendix).

CIBERSORTx analytical tool [33] with the high-resolution mode was used to estimate cell type-specific gene expression for neurons and astrocytes. The analysis requires three types of files: signature matrix file, mixture file and gene subset file. The signature matrix was constructed based on cell type-specific RNA expression data of the mouse cerebral cortex [34]. The mixture files (ACC and DLPFC) consisted of DESeq2-normalized gene expression values for all genes across all samples. DESeq2 uses median-of-ratios method for normalization. The gene subset file contained all Winter model genes. The CIBERSORTx analysis was performed using the analysis module “3. Impute Cell Expression” and analysis type “High-Resolution”. Optional merged class or ground truth files were not used. Batch correction (B-mode) was enabled and quantile normalization was disabled. The window size used for deconvolution was 20. In the end, imputation of cell type-specific expressions did not provide coherent results for all the DEGs, and this problem was present especially with astrocyte-specific expressions. Therefore, a decision was made to use both CIBERSORTx imputed gene expressions and bulk RNA expressions (separately) to perform the brain energy metabolism simulations.

### Description of Winter model and its modification

In this study, we used a biophysically detailed model of brain energy metabolism developed by Winter et al. (2018) [27]. The Winter model consists of five compartments (artery, capillary, extracellular space, neuron and astrocyte) and 64 reactions (40 metabolic reactions and 24 transport reactions). The model describes cytosolic energy metabolism processes – glycolysis and pentose-phosphate pathway (PPP) – in detail, whereas mitochondrial respiration is described by one reaction. In the model, neuronal stimulation is described as a combination of (a) release of glutamate from neurons into the extracellular space coupled with an influx of sodium from the extracellular space to the neurons, and (b) an increase in cerebral blood flow (CBF). A simplified illustration of the Winter model is presented in Fig 1.

**Fig 1.**
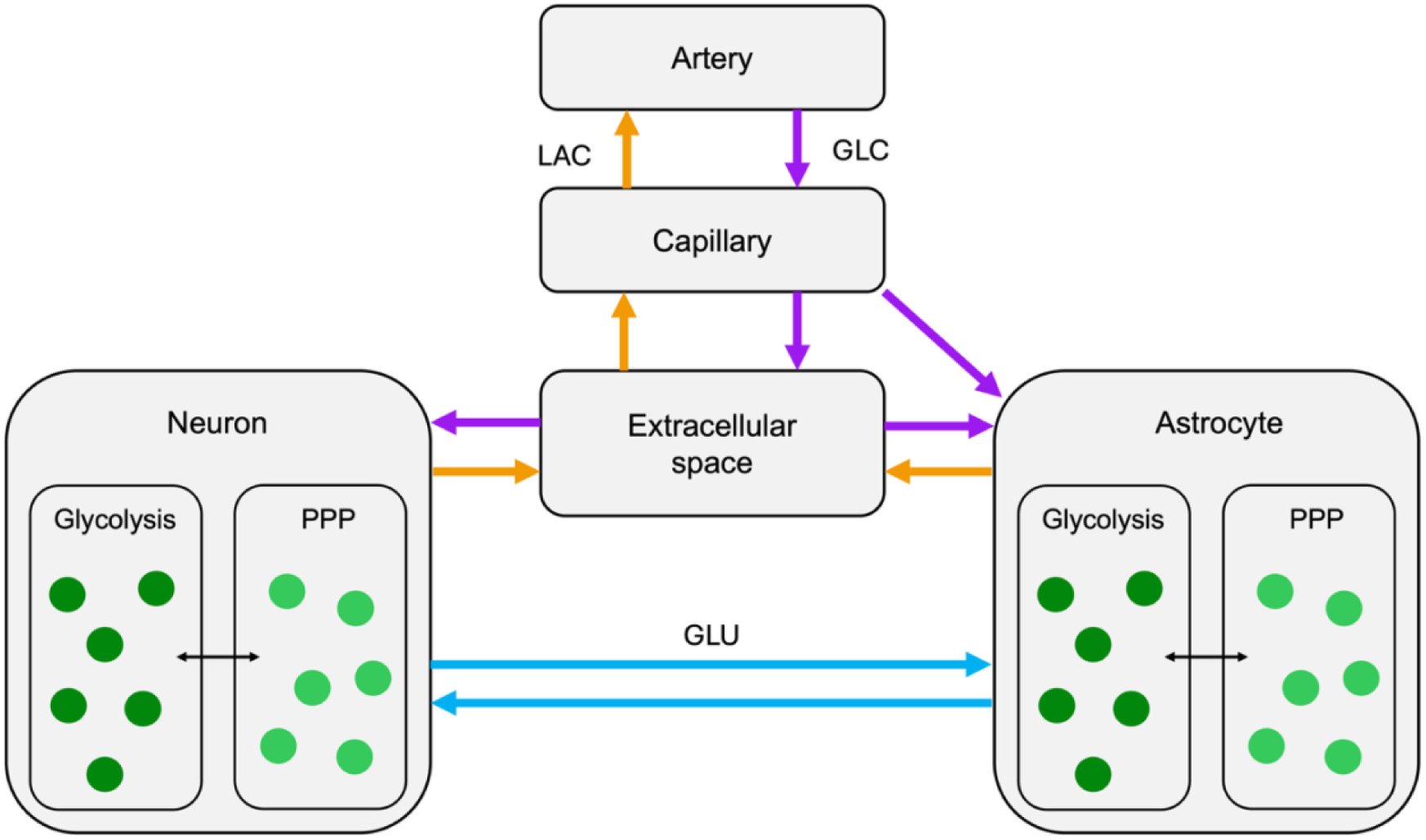
Simplified illustration of the Winter model. Purple and orange arrows indicate the direction of glucose (GLC) and lactate (LAC) flux at steady state, respectively. Blue arrows indicate glutamate (GLU) flux after neuronal stimulation. Dark green circles represent metabolic reactions of glycolysis and light green circles those of pentose-phosphate pathway (PPP). Adapted from [27].

We used the Winter model to study how altered gene expression influences the concentrations of central energy metabolites: ATP, glucose (GLC), lactate (LAC), pyruvate (PYR), and NAD^+^/NADH ratio, in neurons and astrocytes. Before the brain energy metabolism simulations, we made some modifications to the Winter model to improve its numerical stability. We multiplied the “vMITO2 (incl. Volumes)” function with 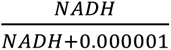 and the “vHK (HS)” function with 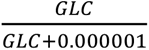. These modifications prevented negative concentrations of NADH and glucose in the subject-specific simulations. Hereafter, we will refer to this modified Winter model as mWinter model. We used the mWinter model for all the simulations. The mWinter model and instructions on how to replicate the analyses presented in this article are provided in ModelDB (https://modeldb.science/2020514, password is ‘SCZmetabolism’).

### Brain energy metabolism simulations in COPASI

The brain energy metabolism simulations were performed using COPASI simulator [35] which uses the Runge-Kutta integration method for simulating ordinary differential equations. The COPASI file of the original Winter model was downloaded from BioModels database [36], where the model is stored with an identifier MODEL1603240000.

The modified mWinter model with default parameters was used as a model of cytosolic energy metabolism in the healthy brain. The schizophrenia simulations were done by adjusting the parameter(s) of the reactions of interest. The reaction parameters were adjusted either based on CIBERSORTx data and bulk data (separately) or solely based on bulk data if CIBERSORTx expressions were not available.

We performed simulations using both population-average and subject-specific data. In the population-average simulations, coefficients were derived by comparing the mean gene expression of SCZ samples with that of HC samples. The coefficients were then used to modify the model parameters separately (single-gene simulations) and collectively (multi-gene simulations). The reactions, modified parameters, and coefficients are listed in S3 Appendix. In the case of subject-specific simulations, the coefficients were calculated for each sample, and all samples were simulated individually as multi-gene simulations.

Subject-specific simulations were also performed with added noise to assess whether the predicted differences between SCZ and HC samples remained significant under small random perturbations in the expression of energy metabolism–associated genes. First, we calculated the coefficient of variation (standard deviation divided by mean) for each DEG across all HC samples to assess the variability of gene expression relative to the mean. We obtained an average coefficient of variation around 0.2. We opted to use a 10 % subject-specific perturbation of the expression of all energy metabolism-associated genes. To do this, we multiplied each model parameter listed in S3 Appendix by a Gaussian random number (independently for each subject) with mean value of 1 and standard deviation of 0.02 in addition to possible alterations of the model parameter according to the gene expression data.

We performed a time course analysis to study how the metabolite concentrations respond to neuronal stimulus over time. Before the energy metabolism simulations, we investigated how the two stimulation components of the mWinter model (glutamate release coupled with an influx of sodium and increased CBF) separately and together influence the metabolite concentrations during the time course. The effects of only glutamate release were studied by setting global parameter delta_F = 0. The effects of only increased CBF were studied by setting global parameters is_stimulated = 0 and stimulus = 0. Other model parameters were not modified. This analysis showed that glutamate release has a larger impact on the time courses of the metabolites than increased CBF (Fig 2).

**Fig 2.**
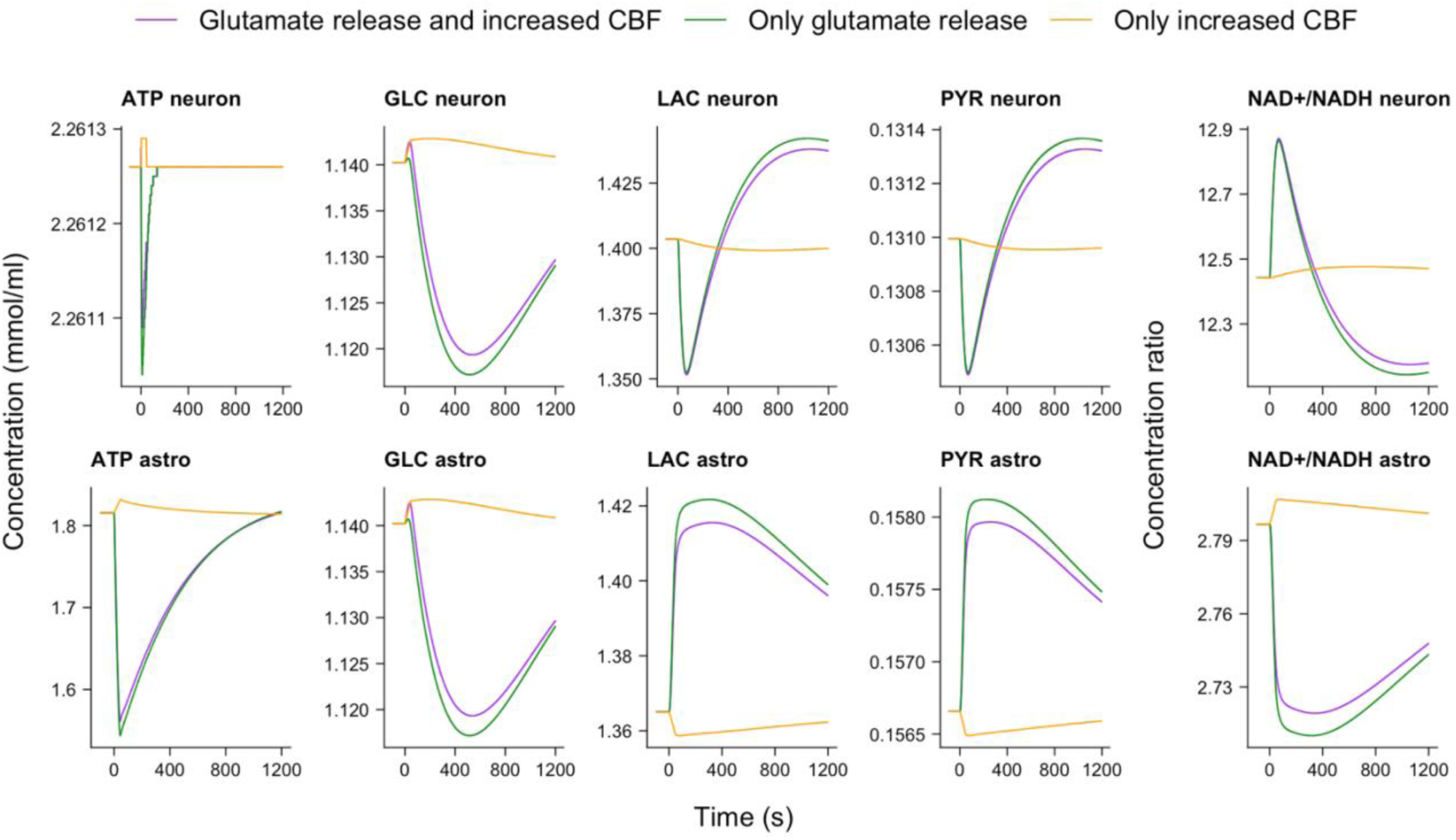
Time course simulations of the mWinter model showing how separated and combined stimuli affect central energy metabolite concentrations. Simulations were run to steady state before stimulation, which was applied at 30,000 s. For clarity, data before 29,900 s are not shown, and time was shifted so that the stimulus appears at 0 s in the plots. The concentrations were recorded at 0.1 s time intervals. An interval size of 0.01 s was also tested, but since no noticeable differences were observed, the larger interval was chosen. Astrocyte (astro), cerebral blood flow (CBF), glucose (GLC), lactate (LAC), pyruvate (PYR).

### Statistical analysis and visualization

We used Wilcoxon signed-rank test to study the statistical significance of each metabolite concentration at baseline in SCZ vs. HC. Benjamini-Hochberg (BH) procedure was used to control the false discovery rate (FDR). In the case of time course plots portraying the difference between baseline and maximum/minimum concentration value, permutation test was used to assess significance. The number of permutations was 10,000. Again, BH procedure was used to control the FDR. All the plots were made in R using ggplot2 (v3.5.1).

## RESULTS

### RNA sequencing data show region-specific altered expression of energy metabolism genes in schizophrenia

Bulk RNA-seq from the ACC and DLPFC was analyzed to investigate alterations in cytosolic energy metabolism in schizophrenia. First, reactions of the Winter model were mapped to their corresponding proteins and genes. Next, DE analysis was performed to determine how the expression of these genes is altered in schizophrenia.

Out of the 85 Winter model genes, three were upregulated and six were downregulated in the ACC (Fig 3A). In the DLPFC, seven genes were upregulated and 17 were downregulated in schizophrenia (Fig 3B). The names of the DEGs, log2 fold changes and adjusted p-values are presented in Table A (ACC) and Table B (DLPFC) in S4 Appendix.

**Fig 3.**
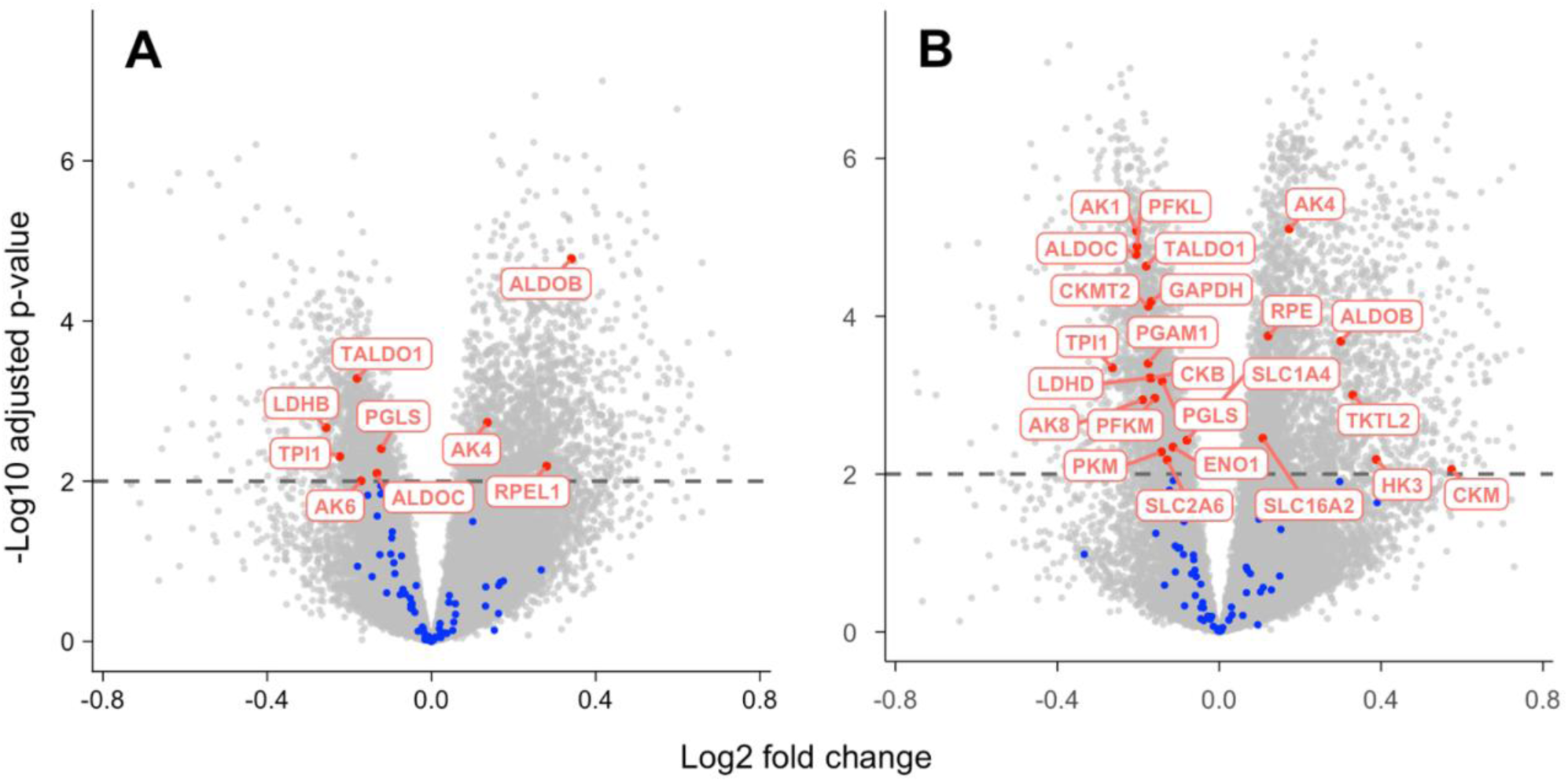
Volcano plots of the differential expression analysis results. (A) Anterior cingulate cortex (ACC). (B) Dorsolateral prefrontal cortex (DLPFC). The gray dots represent all the transcripts in the data sets. The colored dots represent the Winter model genes: the blue ones are non-significant (adj. p-value > 0.01), and the red ones are significant (adj. p-value < 0.01). Due to axis limiting, 25 gray dots were cropped out of the plot in (A) and 50 in (B).

### Simulations based on RNA sequencing data of energy metabolism genes suggest altered baseline concentrations of key metabolites in schizophrenia

After the DE analysis, we adjusted the mWinter model parameters according to the gene expression alterations and simulated how these alterations affect brain energy metabolism in schizophrenia. However, due to the reasons presented in S2 Appendix, some of the DEGs could not be included in the simulations. In the end, we could simulate the effects two DEGs (*LDHB* and *PGLS*) in neurons and one (*PGLS*) in astrocytes based on both bulk and cell type-specific data of the ACC. For the DLPFC, we could simulate the effects of three DEGs (*PFKM*, *PKM*, and *PGLS*) in neurons and one (*PGLS*) in astrocytes using cell type-specific gene expression data. Using bulk data of the DLPFC, we could simulate four DE genes (*PFKM*, *PKM*, *PGLS,* and *RPE)* in neurons and three (*PKM*, *PGLS*, and *CKB*) in astrocytes.

The population-average single-gene simulations suggested that decreased expression of *LDHB* in the ACC and *PFKM* in the DLPFC lead to significant changes in the baseline concentrations of some metabolites in schizophrenia (S5 Appendix). By contrast, altered expression of the other genes (*PGLS* in the ACC, and *PKM*, *PGLS*, *CKB*, and *RPE* in the DLPFC) did not cause significant alterations (difference in baseline concentration > 0.5 %) in any of the metabolite concentrations (S5 Appendix). This result was obtained using both bulk and cell type-specific gene expression data. Consequently, multi-gene simulation of the ACC (altered expression of neuronal *LDHB* and neuronal and astrocytic *PGLS*) yielded the same results as the single-gene simulation of *LDHB*. The same result was observed in the multi-gene simulation of the DLPFC (altered expression of neuronal *PFKM* and *PKM*, and neuronal and astrocytic *PGLS*) vs. the single-gene simulation of *PFKM*. It should be noted that both *LDHB* (encodes the subunits of lactate dehydrogenase 1) and *PFKM* (encodes phosphofructokinase 1) encode enzymes which are the predominant isoenzymes in neurons but not astrocytes [37–39]. Therefore, only the neuronal mWinter model reactions of these two enzymes were modified for the simulations.

Next, we performed subject-specific multi-gene simulations to further assess the effects of altered expression of energy metabolism-related genes in schizophrenia. The results of noiseless subject-specific simulations of the ACC using cell type-specific data showed that there is a significant difference in the baseline concentrations of all studied metabolites in both neurons and astrocytes (Fig A in S6 Appendix). However, when the same simulations were performed with added noise, only neuronal PYR and NAD^+^/NADH ratio had a significant difference between SCZ and HC, whereas the difference was non-significant for the rest of the neuronal and all astrocytic metabolites (Fig B in S6 Appendix). The simulation results revealed that in neurons, the baseline concentration of PYR is decreased and NAD^+^/NADH ratio is increased in the ACC in schizophrenia (Fig 4A). This result was also evident in the simulations using bulk data (Fig C in S6 Appendix).

**Fig 4.**
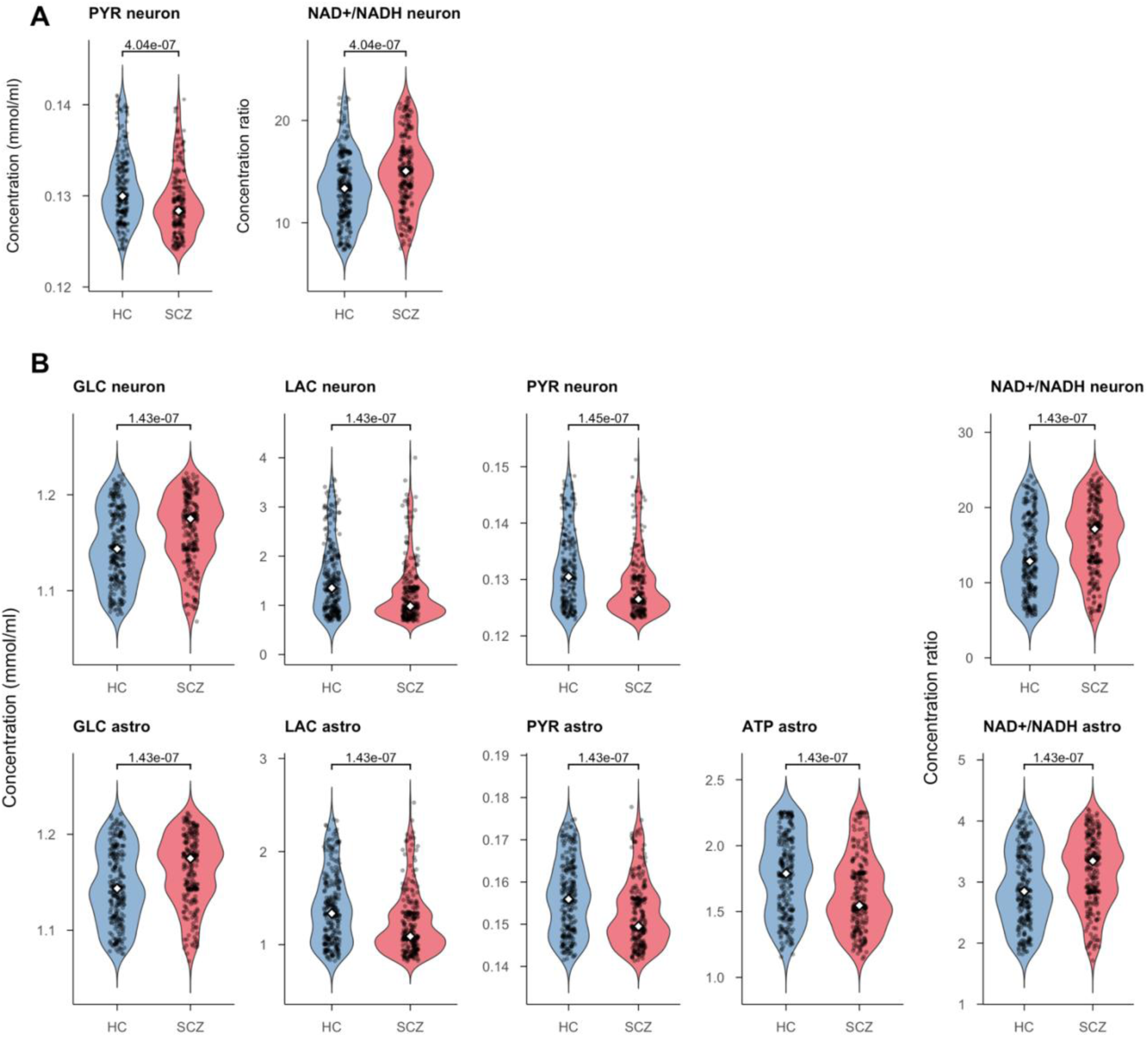
Significant metabolites of the noiseless subject-specific simulations using cell type-specific gene expression data. (A) Anterior cingulate cortex (ACC). (B) Dorsolateral prefrontal cortex (DLPFC). Outliers (values more than three interquartile ranges away from the median in any metabolite, n = 0 for the ACC, n = 71 for the DLPFC) have been filtered out of the plots to improve figure clarity. All samples have been included in median and p-value computation. Adjusted p-values are marked at the top of the plots. The white diamond marks the median of concentrations. Each black dot represents an individual. Healthy control (HC), schizophrenia (SCZ), glucose (GLC), lactate (LAC), pyruvate (PYR), astrocyte (astro).

For the DLPFC, the results of the noiseless subject-specific simulations using cell type-specific data showed that there is a significant difference in the baseline concentrations of all neuronal and astrocytic metabolites in SCZ vs. HC (Fig D in S6 Appendix). When noise was added to the simulations, the concentration differences remained significant for all the metabolites except for neuronal ATP (Fig E in S6 Appendix). The same result was also observed in the simulations using bulk data (Fig F in S6 Appendix). The subject-specific simulations showed higher baseline concentrations of GLC and NAD^+^/NADH ratio, and lower baseline concentrations of LAC and PYR in both neurons and astrocytes, and lower astrocytic ATP, in schizophrenia (Fig 4B).

Taken together, the brain energy metabolism simulations show cell type- and brain region-specific alterations in the baseline concentrations of central energy metabolites in schizophrenia. In the ACC, there are changes in neuronal pyruvate and NAD^+^/NADH ratio, driven by decreased *LDHB* expression. Furthermore, there are significant changes in the baseline concentrations of all neuronal and astrocytic metabolites except for neuronal ATP in the DLPFC, driven by decreased *PFKM* expression.

### Energy metabolism simulations suggest altered time courses of metabolite concentrations in response to neuronal activity in schizophrenia

In addition to assessing the effects of DEGs on the baseline concentrations, we studied how gene expression alterations influence the time courses of metabolite concentrations in response to neuronal stimulus. We assessed the significance of the difference between the baseline and maximum (max) concentration, and the baseline and minimum (min) concentration of the 20 min (1,200 s) time course in SCZ vs. HC. The time courses of only those metabolites which were significantly altered (adj. p-value < 0.05) in both the noisy and noiseless simulations are presented in this section. Violin plots portraying the differences between baseline and max/min in SCZ vs. HC for all metabolites in the ACC and DLPFC are presented in S7 Appendix, with adj. p-values.

The results of the noiseless subject-specific time course simulations of the ACC provided similar results to the baseline simulations: the difference between the baseline and maximum concentration and the baseline and minimum concentration of most metabolites during the time course were significantly altered in SCZ. However, when noise was added to the simulations, only changes in the time courses of neuronal PYR and NAD^+^/NADH ratio remained significant (Fig 5A). More specifically, in the case of PYR, there was a significant difference between the baseline and both the maximum and minimum in SCZ vs. HC. In the case of NAD^+^/NADH ratio, the difference between the maximum concentration and baseline was significantly altered in SCZ, but the difference between minimum concentration and baseline was not.

**Fig 5.**
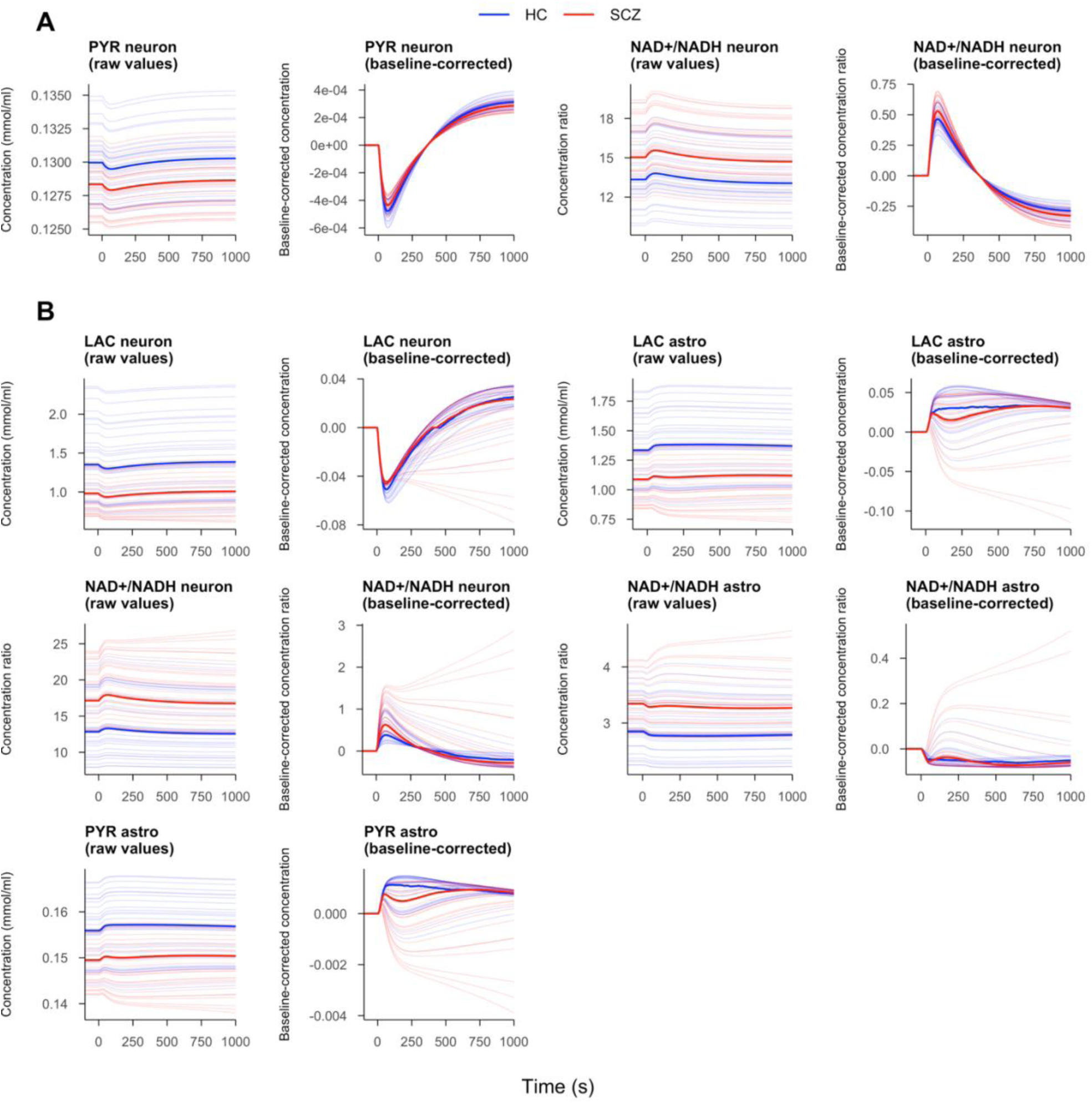
Significant metabolites of the noiseless subject-specific time course simulations, using cell-type specific gene expression data. (A) Anterior cingulate cortex (ACC). (B) Dorsolateral prefrontal cortex (DLPFC). Bold lines represent the median time course of all SCZ vs. HC. First, the subjects with outlier metabolite values (concentrations more than two median absolute deviations away from the median) were filtered out, and then 30 SCZ and 30 HC of the remaining individuals were plotted (thin lines). Simulations were run to steady state before stimulation, which was applied at 30,000 s. For clarity, data before 29,900 s are not shown, and time was shifted so that the stimulus appears at 0 s in the plots. The simulations were run for 1,200 s after the stimulus but only the first 1,000 s are shown because no significant changes occurred thereafter. Healthy control (HC), schizophrenia (SCZ), glucose (GLC), lactate (LAC), pyruvate (PYR), astrocyte (astro).

In the noiseless subject-specific time course simulations of the DLPFC, we observed that the differences between the baseline and maximum concentration of neuronal and astrocytic LAC, astrocytic PYR, and neuronal NAD^+^/NADH ratio were significantly altered in SCZ in both noisy and noiseless simulations. When it came to the differences between the baseline and minimum concentration, there was a significant alteration in neuronal LAC, and neuronal and astrocytic NAD^+^/NADH ratio in SCZ in both simulations. The time courses of the significantly altered metabolites of the DLPFC are presented in Fig 5B.

In summary, most metabolites with altered baseline concentrations in schizophrenia also showed significant changes in their time-course trajectories. The ACC simulations showed that the time courses of neuronal pyruvate and the NAD⁺/NADH ratio were affected by altered gene expression. By contrast, fewer metabolites displayed time course changes than baseline alterations in the DLPFC, limited to neuronal and astrocytic lactate and NAD⁺/NADH ratio, and astrocytic pyruvate.

## DISCUSSION

In this study, we integrated RNA-seq data with computational modeling and found region-specific brain energy metabolism dysregulation in schizophrenia linking DE genes with metabolite alterations. The post-mortem DE analysis results show altered expression of cytosolic energy metabolism-associated genes in SCZ vs. HC, with the majority of genes being downregulated. These gene expression alterations are more predominant in the DLPFC than the ACC. Furthermore, the simulations portraying the effects of gene expression changes on metabolite concentrations also show gene-, cell type-, and region-specific alterations of brain energy metabolism in schizophrenia. These alterations are more numerous in the DLPFC, where metabolite changes are present in both neurons and astrocytes in contrast to the ACC, where metabolite changes are present only in neurons. Together, these findings implicate region-specific dysregulation of brain energy metabolism in SCZ.

### Differential expression analysis results

The aim of the DE analysis was to identify energy metabolism-related genes whose expression is altered in schizophrenia and use this information to simulate the effects of altered gene expression on the concentrations of central metabolites. The results of the DE analysis show that there are almost three times as many DEGs in the DLPFC than the ACC which suggests that cytosolic energy metabolism processes are more dysregulated in the former brain region. Previous studies have also suggested that the frontal cortex is particularly affected in schizophrenia [40], although this finding may be biased by more extensive research focus on this area compared to the ACC. However, around two thirds of the DE genes in both brain regions are downregulated (17 out of 24 in the DLPFC and 6 out of 9 in the ACC) which suggests similarities in the dysregulation of bioenergetics in these two regions. Furthermore, the regions share six DEGs – two of them are upregulated (*AK4* and *ALDOB*) and four are downregulated (*ALDOC*, *TPI1*, *PGLS*, *TALDO1*) in both regions. This overlap is especially significant for the ACC where six out of nine DEGs are also differentially expressed in the DLPFC. In summary, the DE analysis results suggest that there are both region-specific and potentially more general changes in cytosolic energy metabolism-associated gene expression in schizophrenia.

The similarities in gene expression patterns between the ACC and the DLPFC can be assessed by taking a closer look on which biological processes the DEGs are involved in. In both brain regions, nearly half of the DEGs encode glycolytic enzymes and almost all of them are downregulated. In fact, the only two upregulated glycolysis genes (*HK3* and *ALDOB*) are not expressed on the protein level in neurons or glia in the cerebral cortex, according to the Human Protein Atlas [41]. When it comes to the genes that encode proteins related to ATP level regulation (such as adenylate and creatine kinases), there is a clear difference between the two brain regions: there are significantly more DEGs in the DLPFC than the ACC (six vs. one). On the contrary, there are more differentially expressed PPP genes in the ACC than the DLPFC (three vs. two). Lastly, it is interesting that although 15 out of 85 Winter model genes encode glucose and lactate transporters, only two of them (*SLC2A6* and *SLC16A2*) were DE in schizophrenia, both in the DLPFC. Furthermore, *SLC2A6* encodes GLUT6 and *SLC16A2* encodes MCT8 (also known as MCT7), and no evidence of these two proteins being significantly expressed in neurons or astrocytes was found [42,43]. These findings are in line with other studies which have often reported expression changes in cytosolic energy metabolism enzymes and less frequently in metabolite transporters [44].

Most of the DEGs identified in this study have also been identified in at least one other transcriptomic or proteomic study on schizophrenia [44,45]. The simulation results show that there are two genes – *LDHB* in the ACC and *PFKM* in the DLPFC – whose decreased expression cause significant energy metabolite alterations in schizophrenia. One proteomic study has identified decreased protein-level expression of LDHB in the ACC white matter but not in the gray matter [46,47]. Moreover, one proteomic study found an increase in PFKM expression in the DLPFC [48], whereas another study found decreased PFK1 mRNA expression in pyramidal neurons but not astrocytes, and a 16 % decrease in PFK1 activity in the DLPFC [49].

Although our DE analysis results for the most part align with previous studies on schizophrenia transcriptomics and proteomics, the interpretation of the results of these studies remains challenging due to substantial variability and even contradictory findings. A large fraction of the early studies on schizophrenia transcriptomics was performed using microarrays where only a limited set of genes can be studied at a time, and the sample sizes of are small (often under 20 individuals per group) which increases the likelihood of biased results, especially considering the heterogeneity of schizophrenia [50,51]. Additionally, until recently [52,53] almost all transcriptomic data has been bulk data, meaning there is still very limited information on gene expression differences between cell types. Lastly, more comprehensive studies are needed to disentangle the effects of variables such as age, sex, disease progression, and medication status on gene expression in schizophrenia [12]. Thus, methodological improvements such as single cell sequencing with larger sample sizes are required to understand the complex polygenic factors behind brain energy metabolism disturbances in schizophrenia.

### Simulation results: ACC

Both baseline and time course multi-gene simulations of the ACC show changes in neuronal pyruvate concentration and NAD^+^/NADH ratio. However, comparison of single- and multi-gene simulations of the same brain region reveal that these changes are predominantly driven by decreased expression of neuronal *LDHB*. This gene encodes the subunits of the lactate dehydrogenase 1 (LDH-1) which is the primary isoenzyme of neurons. LDH-1 catalyzes the conversion of pyruvate to lactate and back using NAD^+^(H) as a coenzyme. In neurons, the enzyme kinetics of LDH-1 isoenzyme cause the reaction to favor the backward direction (lactate is converted to pyruvate and NAD^+^ is converted to NADH) [54]. LDH-1 has a key role in neuronal energy metabolism, as it converts astrocyte-supplied lactate and NAD^+^ to pyruvate and NADH, ensuring adequate supply of substrates for ATP production during high neuronal activity.

The results of this study indicate that decreased *LDHB* expression reduces baseline levels of pyruvate in the ACC neurons in schizophrenia, meaning there is a diminished supply of substrates for ATP production. Additionally, decreased LDHB expression increases neuronal NAD^+^/NADH ratio, shifting neurons toward a more oxidized state. Interestingly, we did not observe a significant change in lactate concentration, which is one of the most consistent findings of imaging studies on schizophrenia [17,55]. This lack of alteration in lactate concentration suggests that there is a compensatory change in lactate transport instead. In the mWinter model there are only two possible fates for neuronal lactate: conversion to pyruvate or export.

In summary, the ACC simulations reveal that expression alterations of most DEGs had negligible effects on the metabolite concentrations. *LDHB* was the only exception, producing a small yet significant change in pyruvate and a more substantial shift in the NAD⁺/NADH ratio. Although gene expression alterations caused only relatively small changes in the metabolite concentrations, the magnitude of these changes is expected since larger disruptions in brain energy metabolism would lead to more severe outcomes like those in inborn errors of metabolism [56]. Moreover, the larger shift in the NAD⁺/NADH ratio may have broader consequences, influencing not only energy metabolism but also oxidative stress, which has been implicated in multiple schizophrenia studies [57,58]. Lastly, the predicted changes in energy metabolism can also be compensated for or exacerbated by alterations in synaptic plasticity and neuronal excitability [59]. Given the limited insights available from imaging studies on metabolite alterations in the ACC, this computational modeling approach provides essential information on the dynamics of energy metabolite alterations and their connection to transcriptomics in the ACC in schizophrenia.

### Simulation results: DLPFC

The simulation results showed that gene expression changes cause more widespread effects on the metabolite concentrations in the DLPFC than the ACC. The concentrations of neuronal and astrocytic lactate and NAD^+^/NADH ratio and astrocytic pyruvate were significantly altered according to both baseline and time course simulations. Additionally, neuronal and astrocytic glucose, neuronal pyruvate and astrocytic ATP had significantly altered baseline concentrations. Again, comparison of single- and multi-gene simulations of the DLPFC revealed that these changes are predominantly driven by decreased expression of neuronal *PFKM*.

*PFKM* encodes an enzyme called phosphofructokinase 1 (PFK-1) which is the predominant isoenzyme in neurons. It catalyzes an important regulatory step of glycolysis – the transfer of phosphate from ATP to fructose-6-phosphate, producing ADP and fructose-1,6-bisphosphate. PFK-1 is allosterically inhibited by ATP and allosterically activated by AMP, meaning this enzyme can regulate the rate of glycolysis based on the cell’s energy requirements.

According to the DLPFC simulations, decreased *PFKM* expression significantly alters the concentrations of almost all studied metabolites in both neurons and astrocytes. Firstly, the simulation results show higher baseline levels of glucose in both cell types, and this observation is in line with studies reporting hypometabolism of glucose in the frontal cortex in schizophrenia [40]. Secondly, the DLPFC simulations reveal lower baseline concentrations of pyruvate and lactate of comparable magnitude in both neurons and astrocytes. To our knowledge, no *in vivo* imaging studies have directly measured pyruvate concentrations in schizophrenia, whereas lactate levels have been relatively well characterized with most of the studies reporting elevated lactate in schizophrenia [17,55]. In contrast, our results suggest that transcriptomic alterations may counteract this increase by slowing glycolysis, through downregulation of *PFKM* for example. The DLPFC simulations also show decreased baseline concentration of astrocytic ATP and increased NAD^+^/NADH ratio in both cell types. There is evidence of reduced ATP availability and increased oxidative stress in the frontal regions in schizophrenia [58], and our results suggest that downregulation of *PFKM* may contribute to these observations.

These results propose a new link between gene expression and metabolite alterations in the DLPFC in schizophrenia. Moreover, although only the *neuronal* PFK-1 reaction is altered in the simulations, metabolite changes emerge in both neurons and astrocytes, with more prominent effects in astrocytes. This observation highlights the importance of the function and regulation of PFK-1 enzyme and how its malfunction may lead to widespread effects on brain energy metabolism.

### Challenges and future perspectives

Although nine DEGs in the ACC and 24 in the DLPFC were identified in the DE analysis, we were able to model only two and five of them, respectively. There are multiple factors which limited the number of DEGs which could be modeled, including limitations of the Winter model and lack of proteomic data. Firstly, the Winter model does not describe each glycolytic reaction separately but instead groups some of them together, which is why we were unable to simulate the altered gene expression of *ALDOB, ALDOC, ENO1, GAPDH, PGAM1,* and *TPI1.* Another limitation of the Winter model is that it describes only the central cytosolic energy metabolism reactions, leaving out some of the glycolysis steps and all mitochondrial reactions, limiting the number of reactions which can be modified according to gene expression data. Evidence suggests that mitochondrial energy metabolism-related genes may also be dysregulated in schizophrenia [13,19,60,61], and therefore the reactions they mediate should be included in models to obtain a more comprehensive view of transcriptomics-driven energy metabolism alterations in the disorder.

Another limitation of the current study was the lack of cell type-specific gene expression data. Neurons and astrocytes have distinct but complementary energy metabolism profiles [16,62], and therefore it would be beneficial to adjust model parameters and perform metabolic simulations according to cell type-specific data. In conclusion, these limitations showcase the need for more detailed energy metabolism models, and more cell type and isoform-specific information on gene and protein expression in the brain which would help build and refine the models.

In conclusion, the current results show brain region-specific cytosolic energy metabolism-related gene expression alterations in the ACC and DLPFC in schizophrenia. Although most of these alterations did not significantly affect metabolite levels, we observed one gene (*LDHB*) that caused minor changes in neuronal energy metabolism in the ACC, while another gene (*PFKM*) led to more widespread effects on both neuronal and astrocytic bioenergetics in the DLPFC. Our novel integration of human post-mortem RNA-seq data with a brain energy metabolism model demonstrates how altered gene expression impacts energy metabolite levels in schizophrenia. This novel approach enables us to study the effects of either a single gene or an entire set of DEGs on brain energy metabolism, both at the population and individual levels. These findings provide important insights into the specific transcriptomic factors that may underlie energy metabolism disruptions in schizophrenia, advancing our understanding of the neurobiological mechanisms of the disorder. In the future, this approach has the potential to inform targeted therapies aimed at restoring normal energy metabolism in the brain, offering new strategies for personalized treatments in schizophrenia and other neuropsychiatric conditions.

## ACKNOWLEDGEMENTS

CommonMind Consortium data: The data were generated as part of the CommonMind Consortium supported by funding from Takeda Pharmaceuticals Company Limited, F. Hoffman-La Roche Ltd and NIH grants R01MH085542, R01MH093725, P50MH066392, P50MH080405, R01MH097276, RO1-MH-075916, P50M096891, P50MH084053S1, R37MH057881, AG02219, AG05138, MH06692, R01MH110921, R01MH109677, R01MH109897, U01MH103392, and contract HHSN271201300031C through IRP NIMH. Brain tissue for the study was obtained from the following brain bank collections: the Mount Sinai NIH Brain and Tissue Repository, the University of Pennsylvania Alzheimer’s Disease Core Center, the University of Pittsburgh NeuroBioBank and Brain and Tissue Repositories, and the NIMH Human Brain Collection Core. CMC Leadership: Panos Roussos, Joseph Buxbaum, Andrew Chess, Schahram Akbarian, Vahram Haroutunian (Icahn School of Medicine at Mount Sinai), Bernie Devlin, David Lewis (University of Pittsburgh), Raquel Gur, Chang-Gyu Hahn (University of Pennsylvania), Enrico Domenici (University of Trento), Mette A. Peters, Solveig Sieberts (Sage Bionetworks), Thomas Lehner, Stefano Marenco, Barbara K. Lipska (NIMH).

## SUPPORTING INFORMATION

## S1 Appendix. Biochemical reactions, proteins and genes of the Winter model

Metabolic and transport reactions of the Winter model (doi: 10.1177/0271678X17693024), corresponding proteins and genes that encode those proteins. Note that all gene isoforms listed in this table might not be expressed in neurons and/or astrocytes.

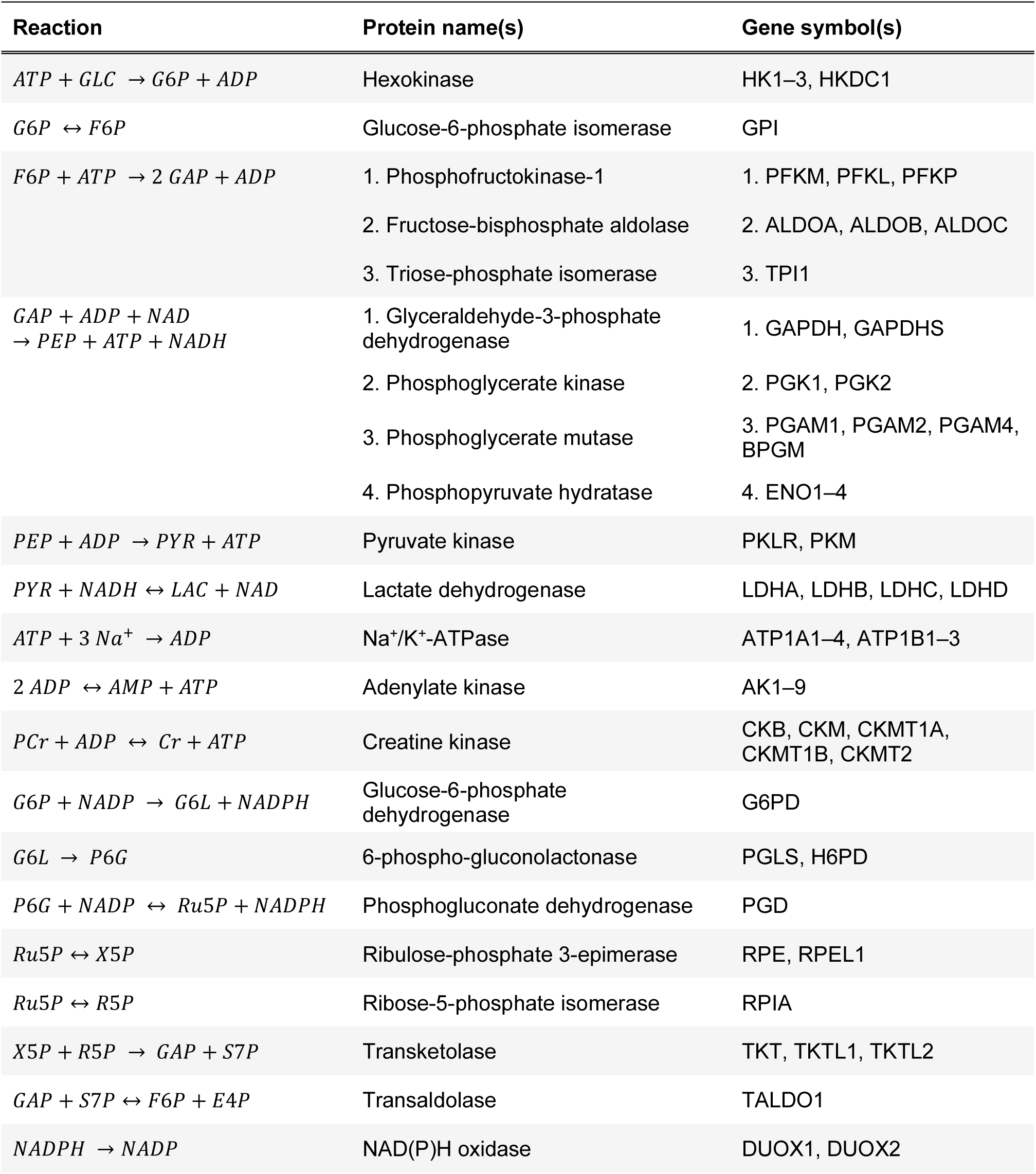
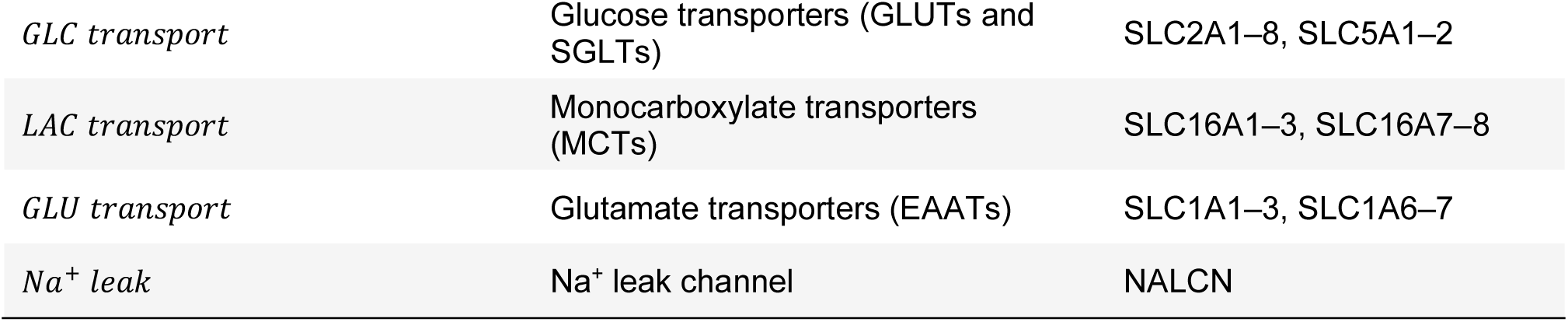

| Reaction | Protein name(s) | Gene symbol(s) |
| --- | --- | --- |
| $ATP + GLC \rightarrow G6P + ADP$ | Hexokinase | HK1–3, HKDC1 |
| $G6P \leftrightarrow F6P$ | Glucose-6-phosphate isomerase | GPI |
| $F6P + ATP \rightarrow 2\ GAP + ADP$ | 1. Phosphofructokinase-1 | 1. PFKM, PFKL, PFKP |
|  | 2. Fructose-bisphosphate aldolase | 2. ALDOA, ALDOB, ALDOC |
|  | 3. Triose-phosphate isomerase | 3. TPI1 |
| $GAP + ADP + NAD \rightarrow PEP + ATP + NADH$ | 1. Glyceraldehyde-3-phosphate dehydrogenase | 1. GAPDH, GAPDHS |
|  | 2. Phosphoglycerate kinase | 2. PGK1, PGK2 |
|  | 3. Phosphoglycerate mutase | 3. PGAM1, PGAM2, PGAM4, BPGM |
|  | 4. Phosphopyruvate hydratase | 4. ENO1–4 |
| $PEP + ADP \rightarrow PYR + ATP$ | Pyruvate kinase | PKLR, PKM |
| $PYR + NADH \leftrightarrow LAC + NAD$ | Lactate dehydrogenase | LDHA, LDHB, LDHC, LDHD |
| $ATP + 3\ Na^+ \rightarrow ADP$ | $Na^+/K^+$-ATPase | ATP1A1–4, ATP1B1–3 |
| $2\ ADP \leftrightarrow AMP + ATP$ | Adenylate kinase | AK1–9 |
| $PCr + ADP \leftrightarrow Cr + ATP$ | Creatine kinase | CKB, CKM, CKMT1A, CKMT1B, CKMT2 |
| $G6P + NADP \rightarrow G6L + NADPH$ | Glucose-6-phosphate dehydrogenase | G6PD |
| $G6L \rightarrow P6G$ | 6-phospho-gluconolactonase | PGLS, H6PD |
| $P6G + NADP \leftrightarrow Ru5P + NADPH$ | Phosphogluconate dehydrogenase | PGD |
| $Ru5P \leftrightarrow X5P$ | Ribulose-phosphate 3-epimerase | RPE, RPEL1 |
| $Ru5P \leftrightarrow R5P$ | Ribose-5-phosphate isomerase | RPIA |
| $X5P + R5P \rightarrow GAP + S7P$ | Transketolase | TKT, TKTL1, TKTL2 |
| $GAP + S7P \leftrightarrow F6P + E4P$ | Transaldolase | TALDO1 |
| $NADPH \rightarrow NADP$ | NAD(P)H oxidase | DUOX1, DUOX2 |
| *GLC transport* | Glucose transporters (GLUTs and SGLTs) | SLC2A1–8, SLC5A1–2 |
| *LAC transport* | Monocarboxylate transporters (MCTs) | SLC16A1–3, SLC16A7–8 |
| *GLU transport* | Glutamate transporters (EAATs) | SLC1A1–3, SLC1A6–7 |
| *Na<sup>+</sup> leak* | Na<sup>+</sup> leak channel | NALCN |

### Abbreviations

ADP: Adenosine diphosphate
AMP: Adenosine monophosphate
ATP: Adenosine triphosphate
(P)Cr: (Phospho)creatine
E4P: Erythrose-4-phosphate
F6P: Fructose-6-phosphate
GAP: Glyceraldehyde-3-phosphate
GLC: Glucose
GLU: Glutamate
G6L: 6-phosphogluconolactone
G6P: Glucose-6-phosphate
LAC: Lactate
NAD(H): Nicotinamide adenine dinucleotide
NADP(H): Nicotinamide adenine dinucleotide phosphate
PEP: Phosphoenolpyruvate
PYR: Pyruvate
P6G: 6-phosphogluconate
Ru5P: Ribulose-5-phosphate
R5P: Ribose-5-phosphate
S7P: Sedoheptulose-7-phosphate
X5P: Xylulose-5-phosphate

## S2 Appendix. Protein-level expressions and selection process of differentially expressed genes for modeling

Multiple differentially expressed genes had to be removed before the simulations, because they could not be modelled by the Winter model. In a few instances, the model combines several metabolic reactions into one model reaction, but these reactions are characterized by only one of the involved proteins (see S1 Appendix). As a result, expression alterations of *ALDOB, ALDOC, ENO1, GAPDH, PGAM1,* and *TPI1* could not be modelled, because the Winter model does not have parameters for these genes. Adenylate kinase genes (*AK1, AK4, AK6,* and *AK8*) were also eliminated: AKs catalyze both the forward and backward reactions, and some of the genes have increased and other decreased expression. Therefore, modulating the AK reaction based on the DE results would not yield coherent results. Lastly, other genes (*CKM, CKMT2, HK3, LDHD, PFKL, RPEL1, SLC2A6, SLC16A2, TALDO1, and TKTL2*) had to be eliminated because we could not find sufficient evidence of protein level expression of these gene in neurons and/or astrocytes (see table below).

Protein expressions of differentially expressed genes in neurons and astrocytes. Note that The Human Protein Atlas [1] presents glial expression, not specifically astrocytic expression.

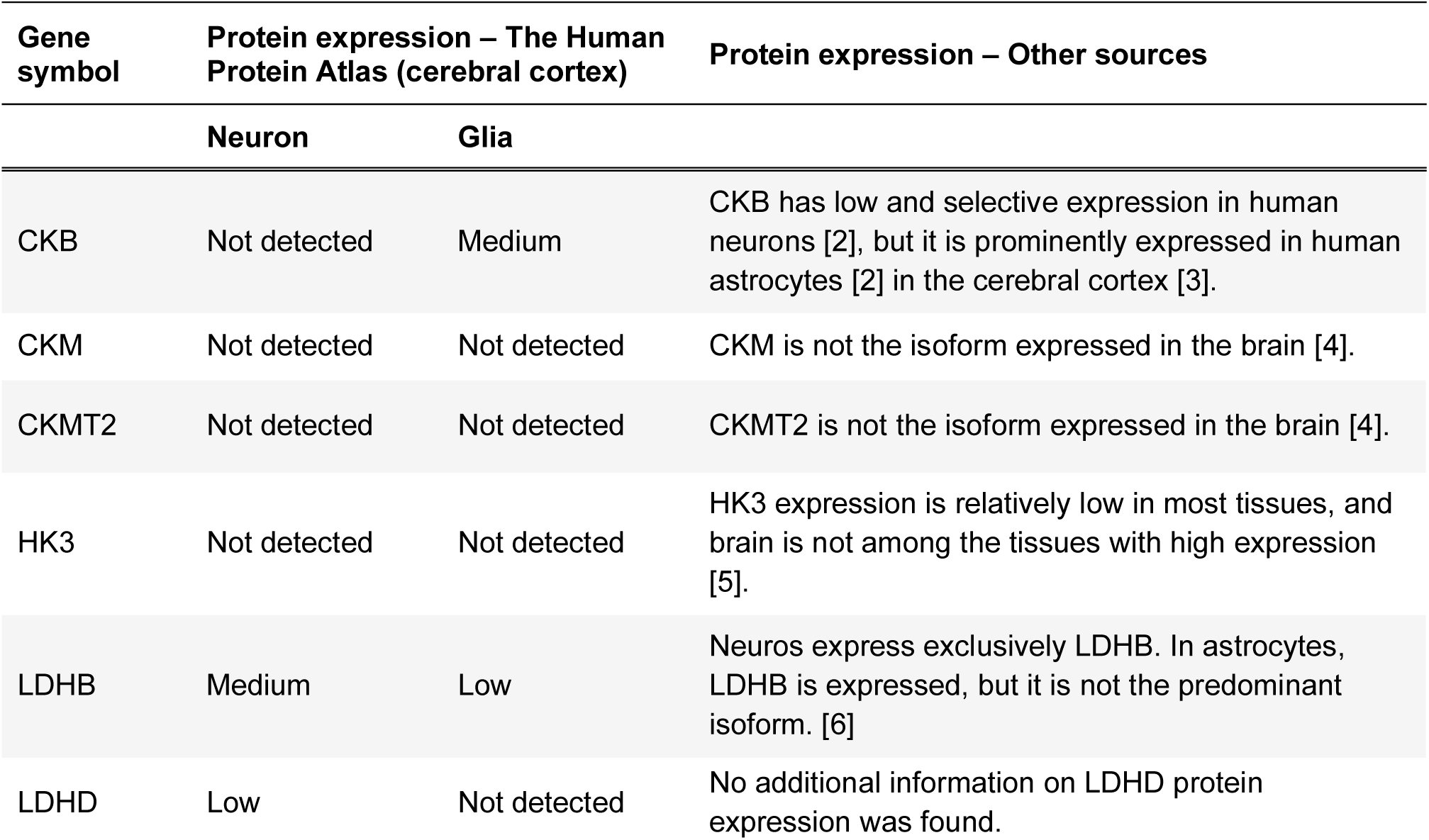
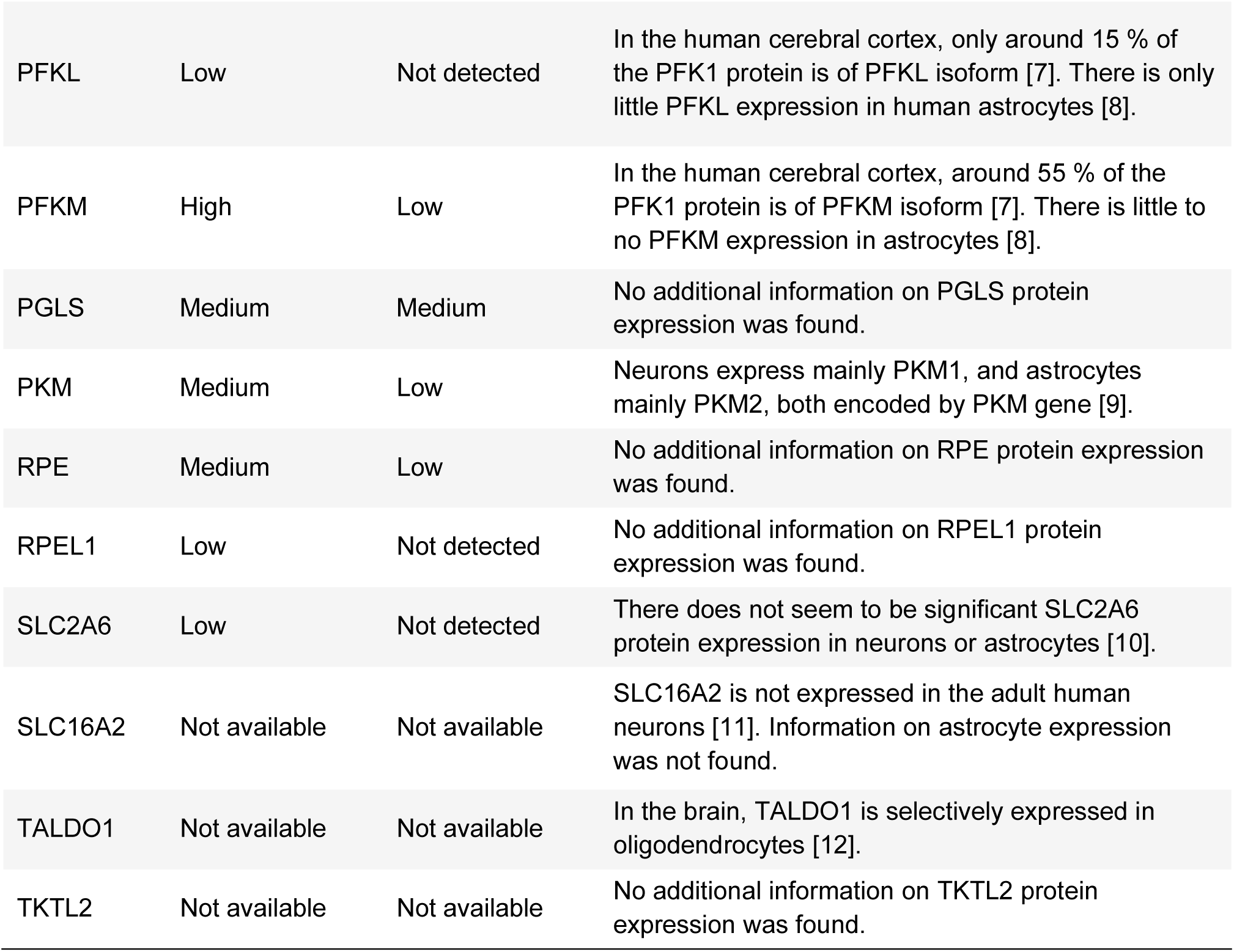

| Gene symbol | Protein expression – The Human Protein Atlas (cerebral cortex) |  | Protein expression – Other sources |
| --- | --- | --- | --- |
|  | Neuron | Glia |  |
| CKB | Not detected | Medium | CKB has low and selective expression in human neurons [2], but it is prominently expressed in human astrocytes [2] in the cerebral cortex [3]. |
| CKM | Not detected | Not detected | CKM is not the isoform expressed in the brain [4]. |
| CKMT2 | Not detected | Not detected | CKMT2 is not the isoform expressed in the brain [4]. |
| HK3 | Not detected | Not detected | HK3 expression is relatively low in most tissues, and brain is not among the tissues with high expression [5]. |
| LDHB | Medium | Low | Neuros express exclusively LDHB. In astrocytes, LDHB is expressed, but it is not the predominant isoform. [6] |
| LDHD | Low | Not detected | No additional information on LDHD protein expression was found. |
| PFKL | Low | Not detected | In the human cerebral cortex, only around 15 % of the PFK1 protein is of PFKL isoform [7]. There is only little PFKL expression in human astrocytes [8]. |
| PFKM | High | Low | In the human cerebral cortex, around 55 % of the PFK1 protein is of PFKM isoform [7]. There is little to no PFKM expression in astrocytes [8]. |
| PGLS | Medium | Medium | No additional information on PGLS protein expression was found. |
| PKM | Medium | Low | Neurons express mainly PKM1, and astrocytes mainly PKM2, both encoded by PKM gene [9]. |
| RPE | Medium | Low | No additional information on RPE protein expression was found. |
| RPEL1 | Low | Not detected | No additional information on RPEL1 protein expression was found. |
| SLC2A6 | Low | Not detected | There does not seem to be significant SLC2A6 protein expression in neurons or astrocytes [10]. |
| SLC16A2 | Not available | Not available | SLC16A2 is not expressed in the adult human neurons [11]. Information on astrocyte expression was not found. |
| TALDO1 | Not available | Not available | In the brain, TALDO1 is selectively expressed in oligodendrocytes [12]. |
| TKTL2 | Not available | Not available | No additional information on TKTL2 protein expression was found. |

---

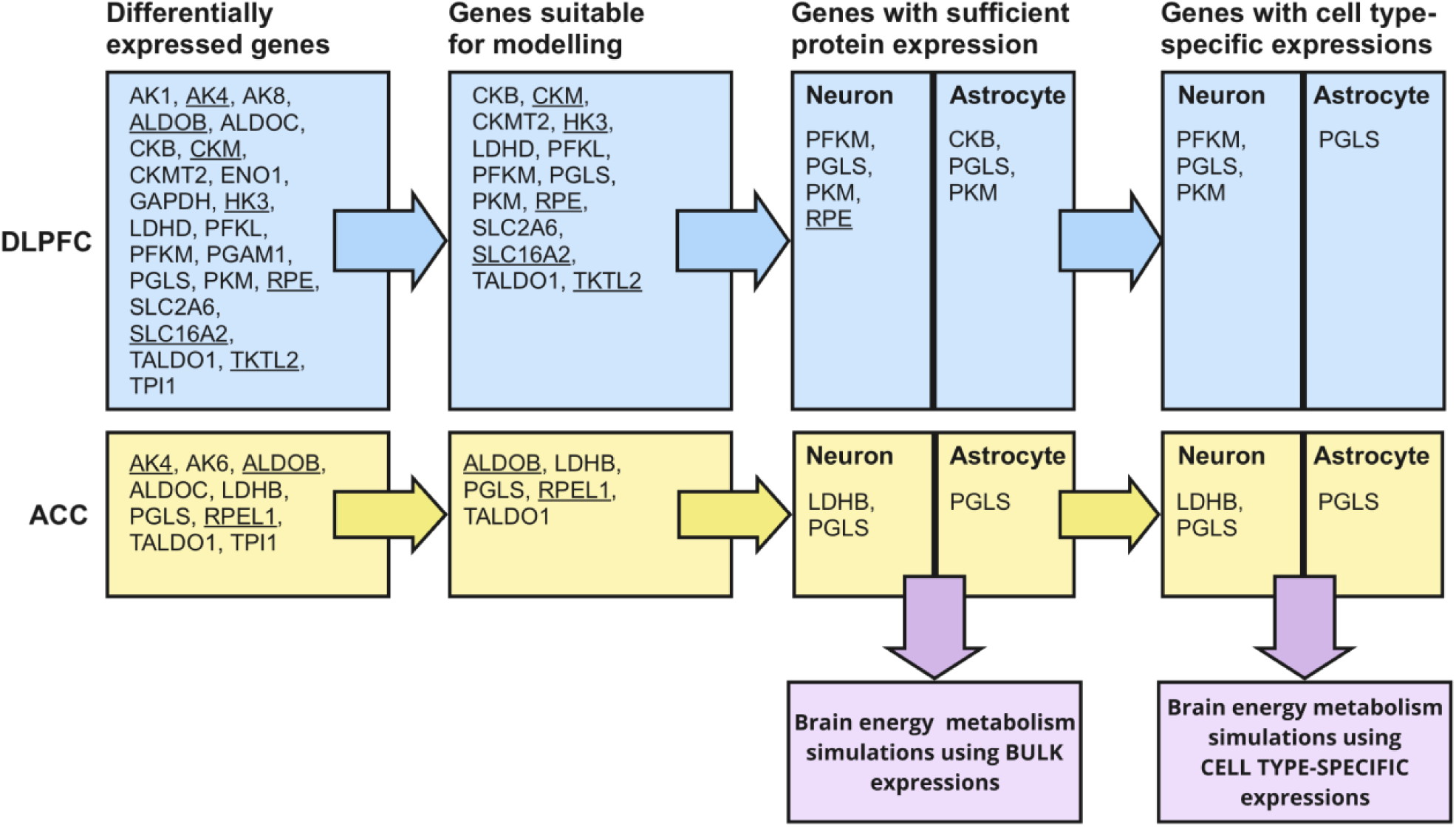

**Diagram of the selection process of DE genes for modeling.** Underlined genes are upregulated, and the rest are downregulated in schizophrenia.

## S3 Appendix. Model reactions and parameters, and multipliers which were used to adjust them

Winter model reactions and parameters which were modified for the simulations. The multiplier column represents the mean expression in schizophrenia subjects (SCZ) vs healthy control (HC), and these values were used to adjust the model parameters. The bulk expressions are DESeq2 normalized (median-of-ratios method), and the cell-type specific expressions are FPKM normalized.

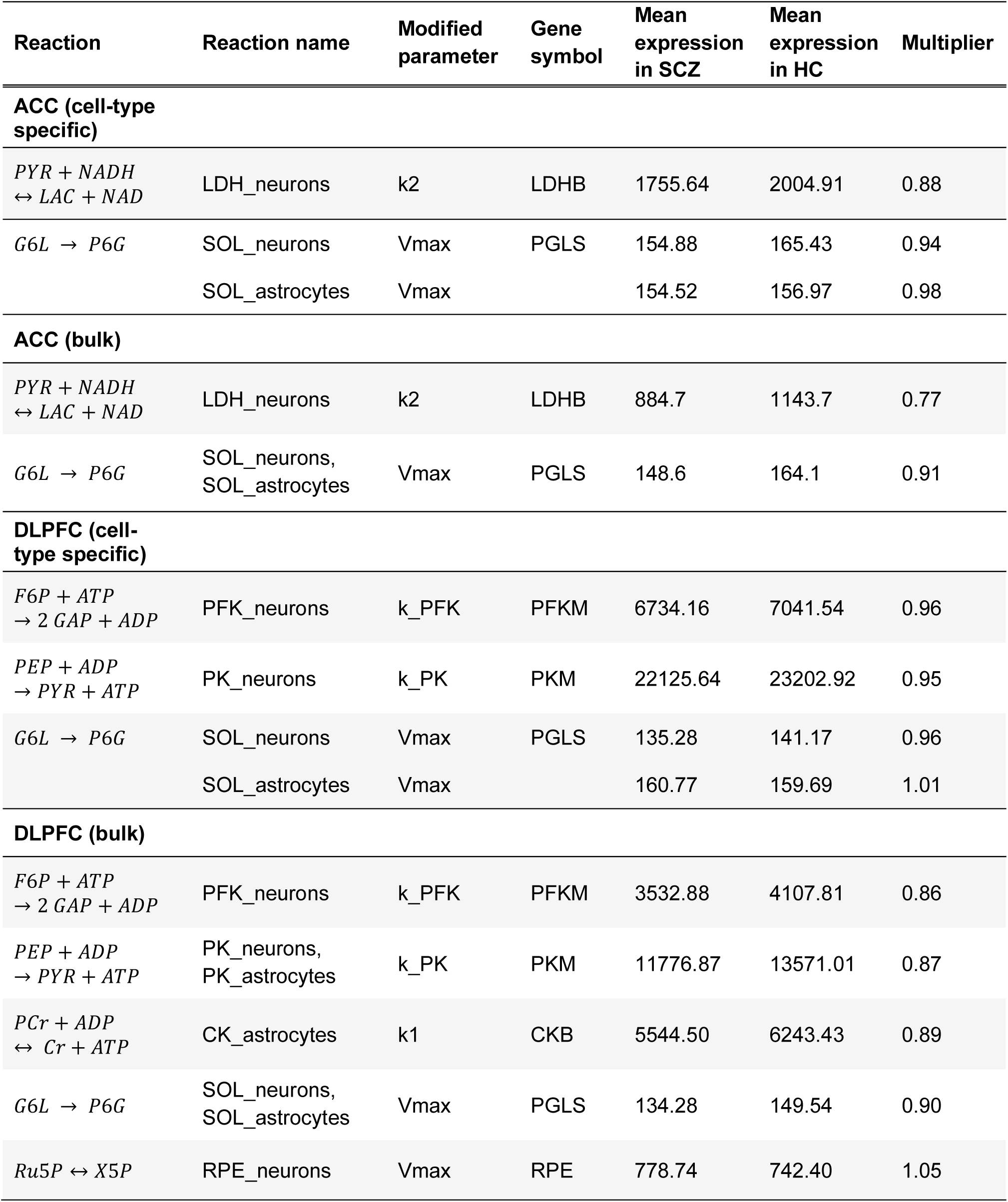

## S4 Appendix. Differentially expressed genes, log2 fold changes and adjusted p-values

**Table A.** Differentially expressed Winter model genes in the **anterior cingulate cortex (ACC).**

| Gene symbol | Log2 fold change (95 % confidence interval) | Adjusted p-value |
| --- | --- | --- |
| ALDOB | 0.34 (0.21, 0.46) | 1.67e-5 |
| TALDO1 | -0.18 (-0.26, -0.10) | 5.21e-4 |
| AK4 | 0.14 (0.07, 0.21) | 1.85e-3 |
| LDHB | -0.25 (-0.39, -0.12) | 2.14e-3 |
| PGLS | -0.12 (-0.19, -0.06) | 3.92e-3 |
| TPI1 | -0.22 (-0.35, -0.10) | 4.93e-3 |
| RPEL1 | 0.28 (0.12, 0.44) | 6.50e-3 |
| ALDOC | -0.13 (-0.21, -0.05) | 7.98e-3 |
| AK6 | -0.17 (-0.28, -0.07) | 9.91e-3 |

**Table B.** Differentially expressed Winter model genes in the **dorsolateral prefrontal cortex (DLPFC)**.

| Gene symbol | Log2 fold change (95 % confidence interval) | Adjusted p-value |
| --- | --- | --- |
| AK4 | 0.17 (0.11, 0.24) | 7.82e-6 |
| AK1 | -0.20 (-0.28, -0.13) | 8.31e-6 |
| PFKL | -0.20 (-0.28, -0.13) | 1.31e-5 |
| ALDOC | -0.21 (-0.28, -0.13) | 1.64e-5 |
| TALDO1 | -0.18 (-0.25, -0.11) | 2.32e-5 |
| GAPDH | -0.17 (-0.24, -0.10) | 6.48e-5 |
| CKMT2 | -0.18 (-0.25, -0.10) | 7.44e-5 |
| RPE | 0.12 (0.07, 0.17) | 1.77e-4 |
| ALDOB | 0.30 (0.16, 0.44) | 2.06e-4 |
| PGAM1 | -0.18 (-0.26, -0.09) | 3.97e-4 |
| TPI1 | -0.26 (-0.40, -0.14) | 4.49e-4 |
| LDHD | -0.17 (-0.26, -0.09) | 6.05e-4 |
| CKB | -0.17 (-0.25, -0.09) | 6.09e-4 |
| PGLS | -0.14 (-0.21, -0.07) | 6.65e-4 |
| TKTL2 | 0.33 (0.16, 0.50) | 9.95e-4 |
| PFKM | -0.16 (-0.24, -0.08) | 1.09e-3 |
| AK8 | -0.19 (-0.29, -0.09) | 1.14e-3 |
| SLC16A2 | 0.11 (0.05, 0.17) | 3.51e-3 |
| SLC1A4 | -0.08 (-0.13, -0.03) | 3.73e-3 |
| ENO1 | -0.11 (-0.18, -0.05) | 4.54e-3 |
| PKM | -0.14 (-0.23, -0.06) | 5.24e-3 |
| HK3 | 0.39 (0.15, 0.63) | 6.52e-3 |
| SLC2A6 | -0.13 (-0.21, -0.05) | 6.60e-3 |
| CKM | 0.57 (0.21, 0.94) | 8.78e-3 |

## S5 Appendix. The results of population-average single-reaction simulations

The colors represent the change in baseline concentration in SCZ vs HC: No difference in concentration (red), insignificant (< 0.5 %) difference in concentration (yellow), significant difference in concentration (green). Plus (+) signs indicate increased concentration, and minus (–) signs decreased concentration in SCZ vs HC. Glucose (GLC), lactate (LAC), pyruvate (PYR), astrocyte (astro).

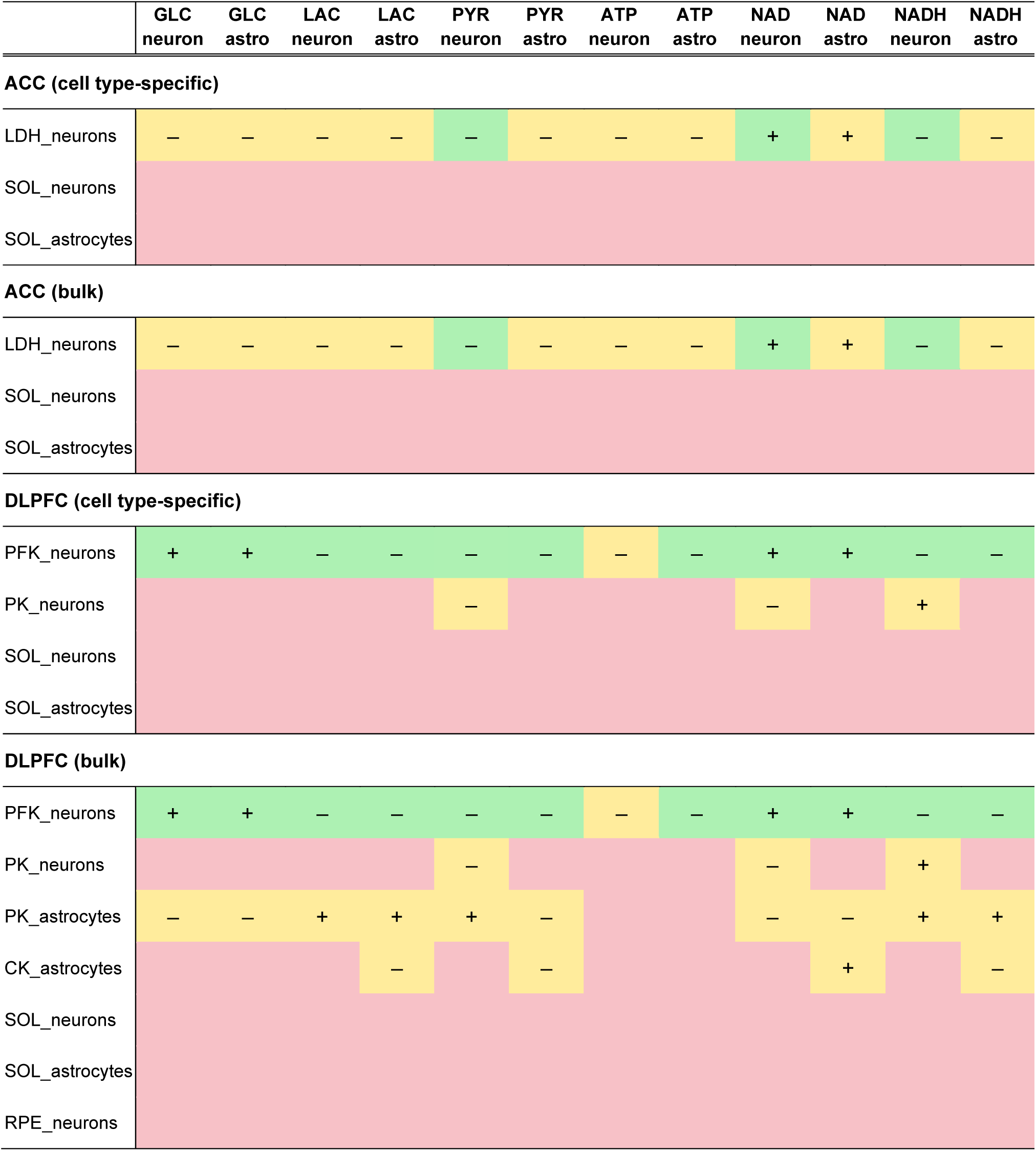

## S6 Appendix. Results of subject-specific multi-gene simulations, effects on the baseline concentrations of metabolites

The p-values indicated above each plot were calculated using Wilcoxon signed-rank test and adjusted using the Benjamini-Hochberg method. In some cases (see figure captions for details) outliers were removed from the figures to improve their clarity. Outliers were defined as values more than three interquartile ranges away from the median (marked with a white diamond). Although outliers were filtered out from the plots, all subjects were included in the statistical tests.

All simulations were run with and without noise. Noise was added by calculating the coefficient of variation (standard deviation divided by mean) for each DEG across all HC samples to assess the variability of gene expression relative to the mean. We obtained an average coefficient of variation around 0.2. We opted to use a 10 % subject-specific perturbation of the expression of all energy metabolism-associated genes. To do this, we multiplied each model parameter listed in S3 Appendix by a Gaussian random number (independently for each subject) with mean value of 1 and standard deviation of 0.02 in addition to possible alterations of the model parameter according to the gene expression data.

**Fig A.**
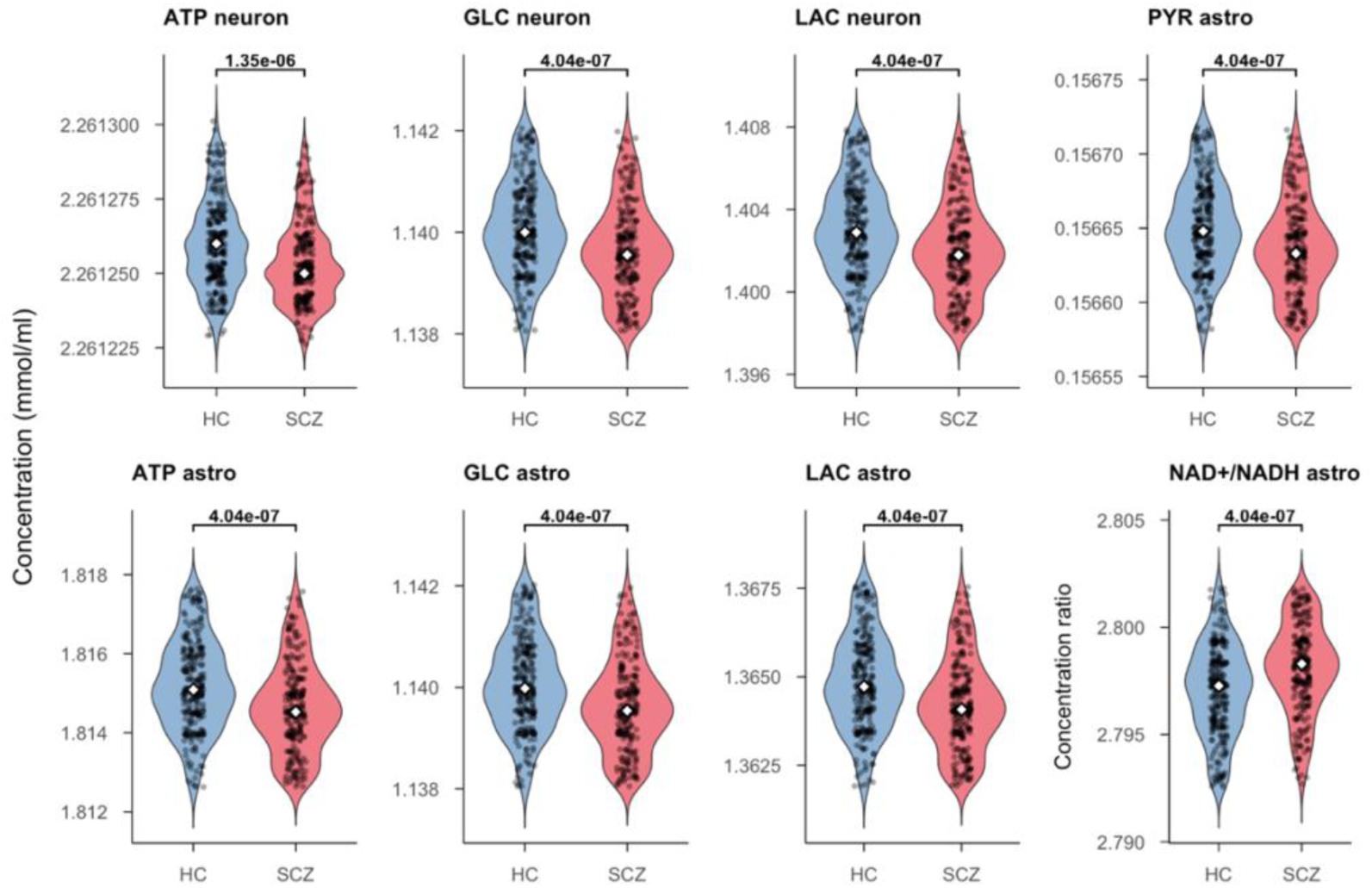
Simulation results of the ACC using cell type-specific data without noise.

**Fig B.**
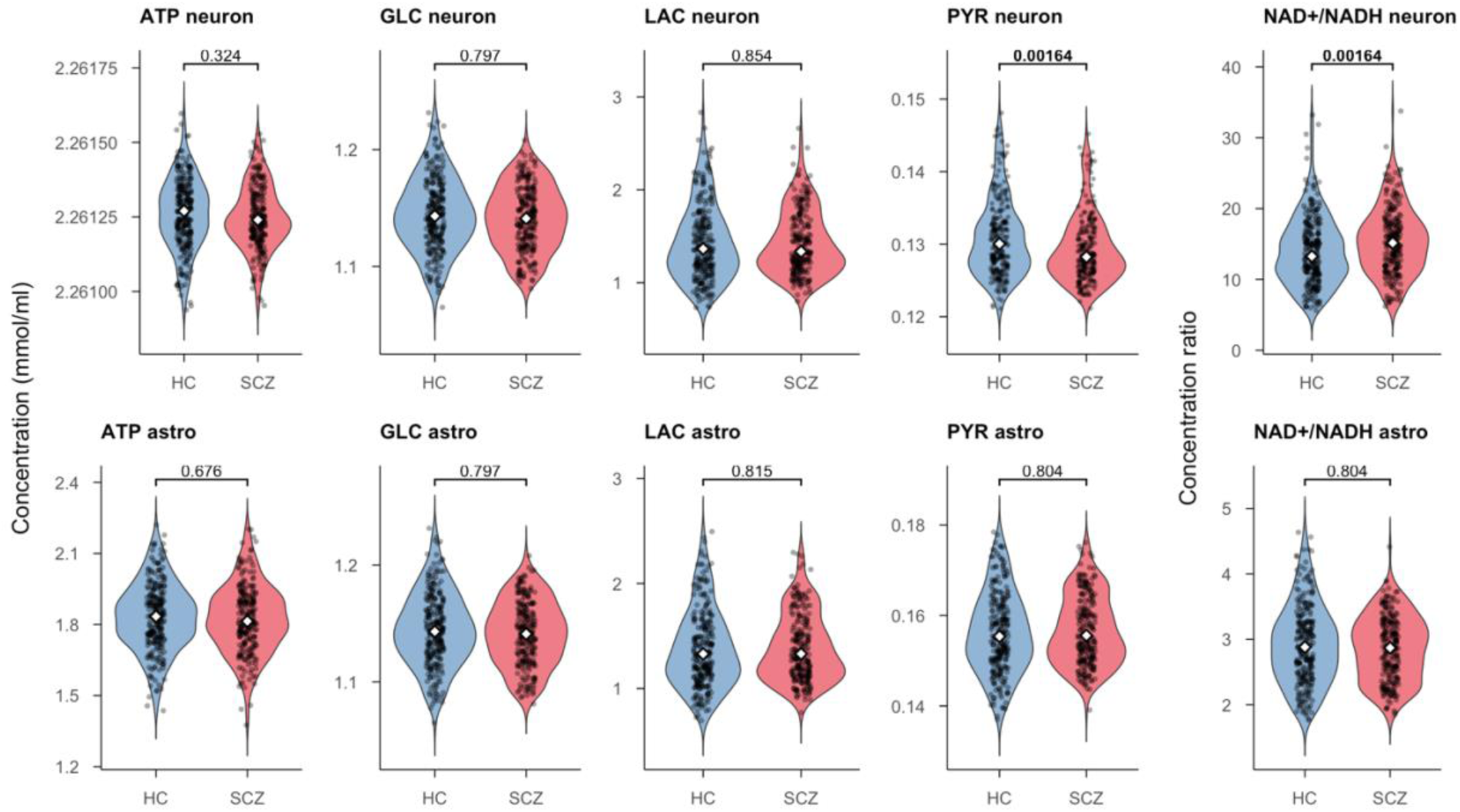
Simulation results of the ACC using cell type-specific data with added noise. Outliers (n = 9) have been removed from the plot.

**Fig C.**
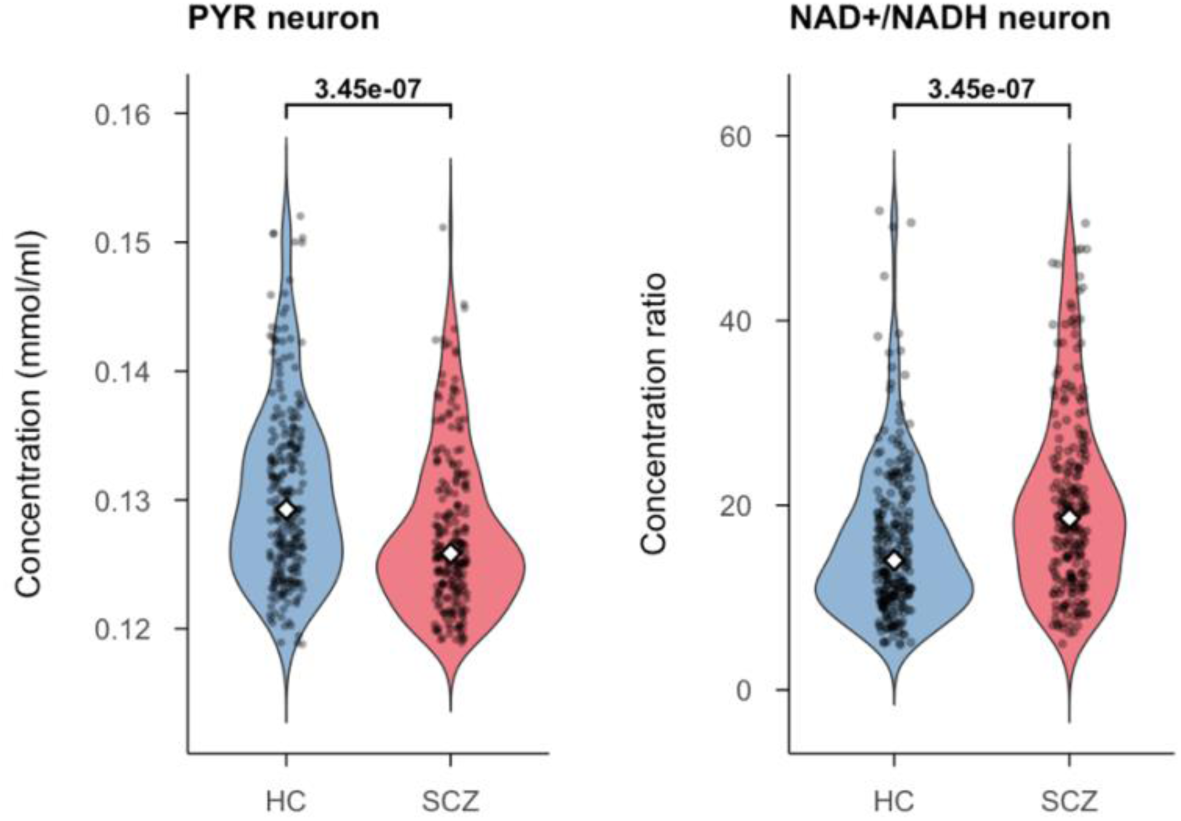
Simulation results of the significant metabolites of the ACC using bulk data without noise. Outliers have been removed from the plot (n = 5).

**Fig D.**
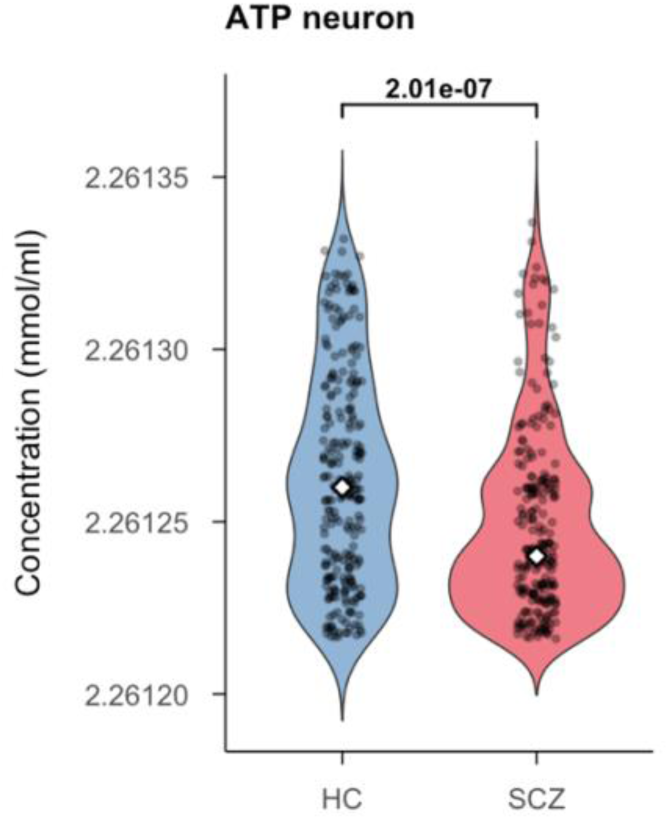
Simulation results of the DLPFC using cell type-specific data without noise. Outliers have been removed from the plot (n = 71).

**Fig E.**
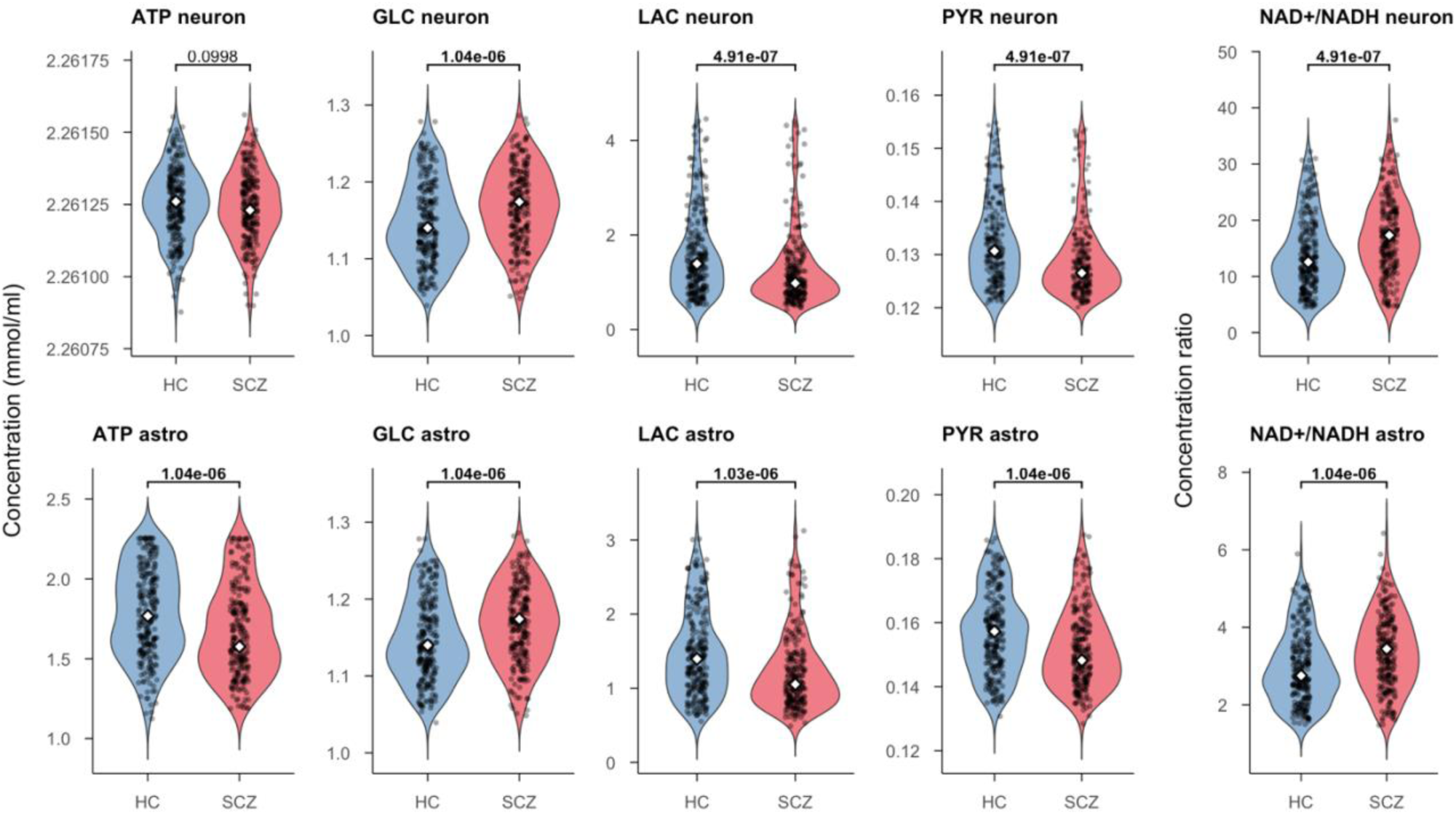
Simulation results of the DLPFC using cell type specific data with added noise. Outliers have been removed from the plot (n = 96).

**Fig F.**
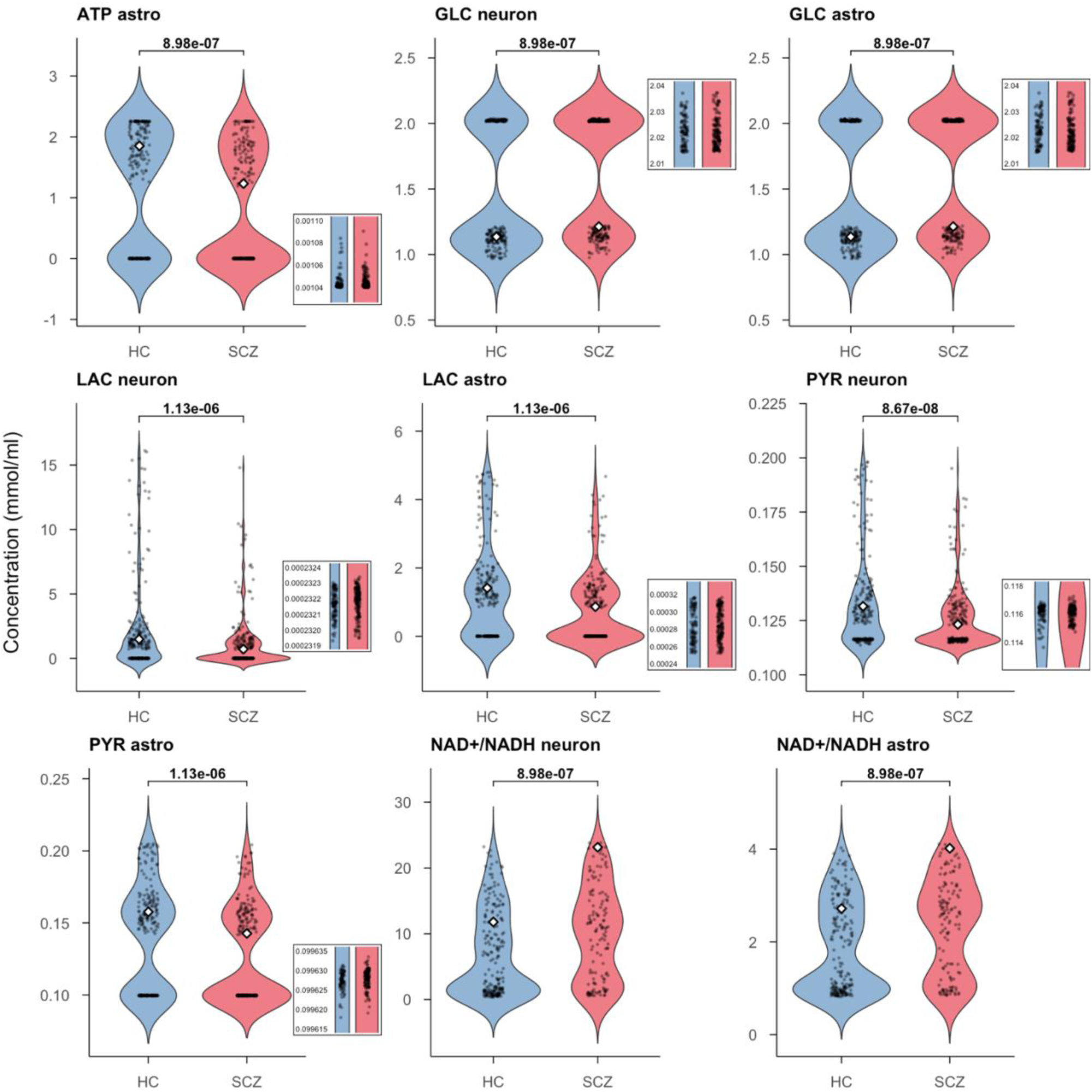
Simulation results of the DLPFC, using bulk data without noise. Outliers have been removed from the plot (n = 117). In the case of NAD^+^/NADH ratio, modified Z score was used to remove the outliers; if |Modified Z| > 3.5, the value was regarded as an outlier (n = 222).

## S7 Appendix. Results of subject-specific multi-gene simulations portraying the difference between baseline and min/max of metabolite concentrations during a time course

The p-values indicated above each plot were calculated using a permutation test with 10,000 permutations and adjusted using the Benjamini-Hochberg method. The metabolites with significant (< 0.05) adj. p-values both with and without added noise are highlighted in gray. In some cases (see figure captions for details) outliers were removed from the figures to improve their clarity. Outliers were defined as values more than three interquartile ranges away from the median (marked with a white diamond). Although outliers were filtered out from the plots, all subjects were included in the statistical tests.

**Fig A.**
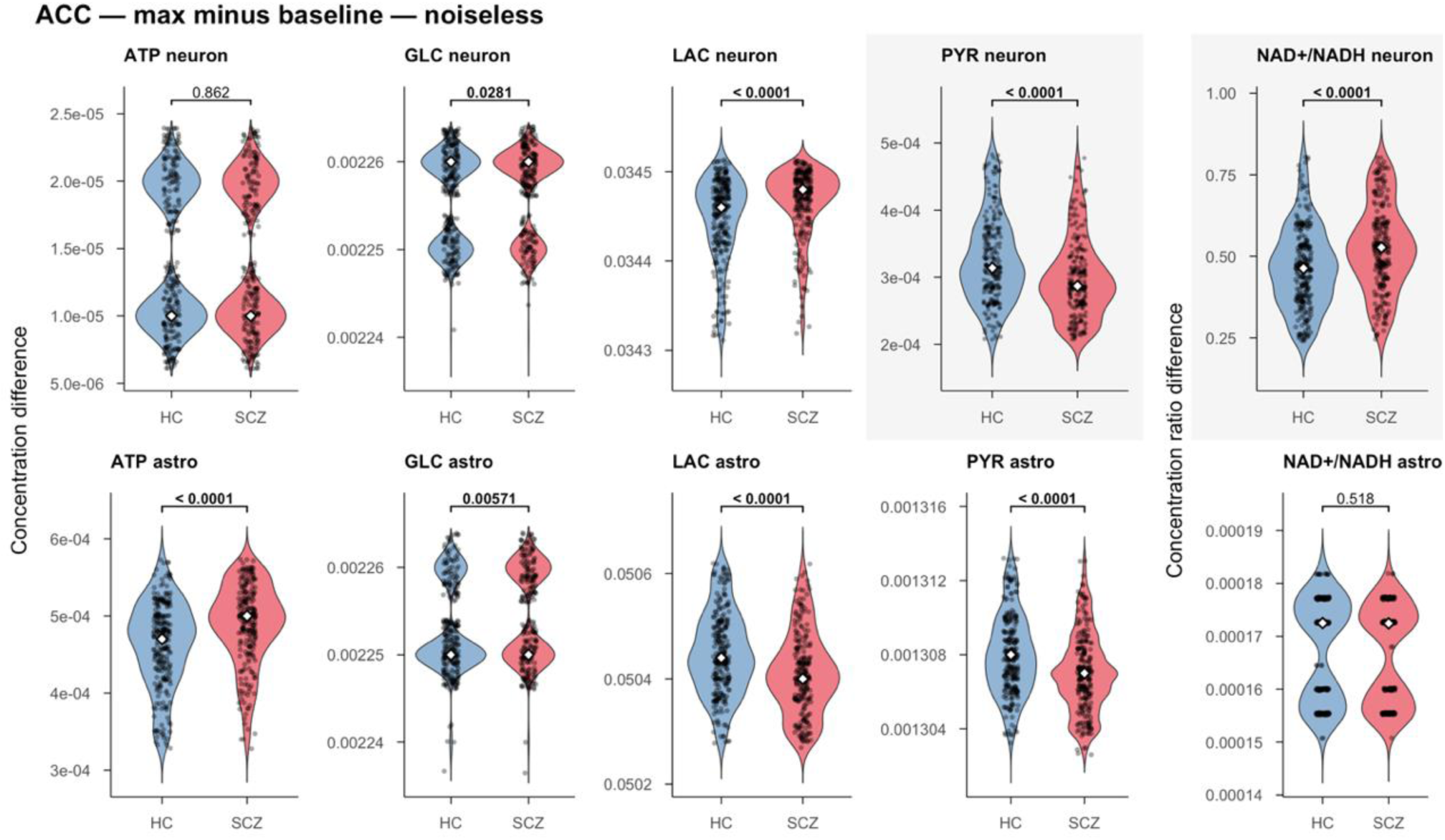
The difference between baseline and maximum concentration of metabolites in the ACC during the 1200 s time course using noiseless data.

**Fig B.**
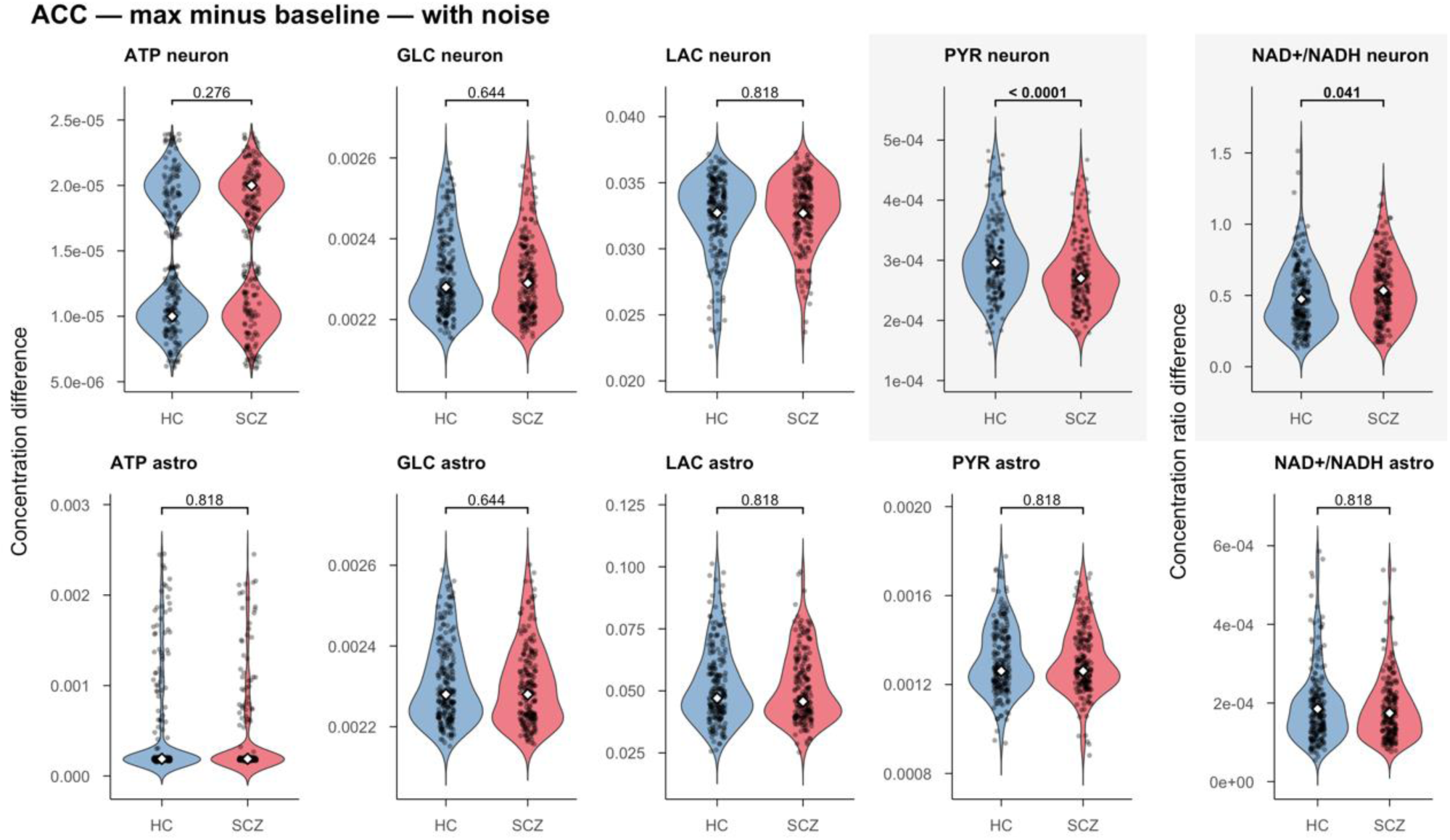
The difference between baseline and maximum concentration of metabolites in the ACC during the 1200 s time course with added noise. Outliers (n = 77) have been removed from the plots.

**Fig C.**
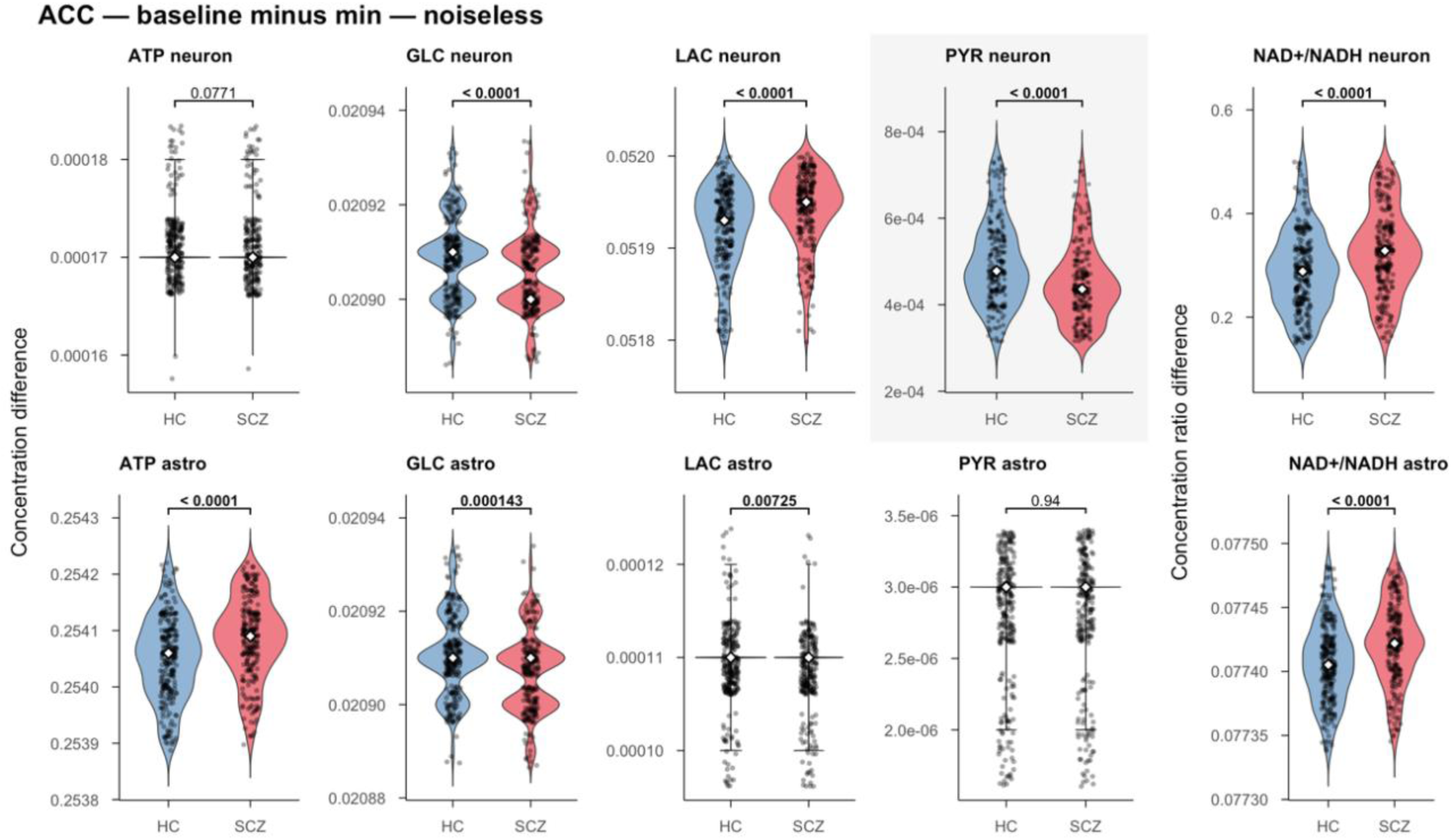
The difference between baseline and minimum concentration of metabolites in the ACC during the 1200 s time course using noiseless data.

**Fig D.**
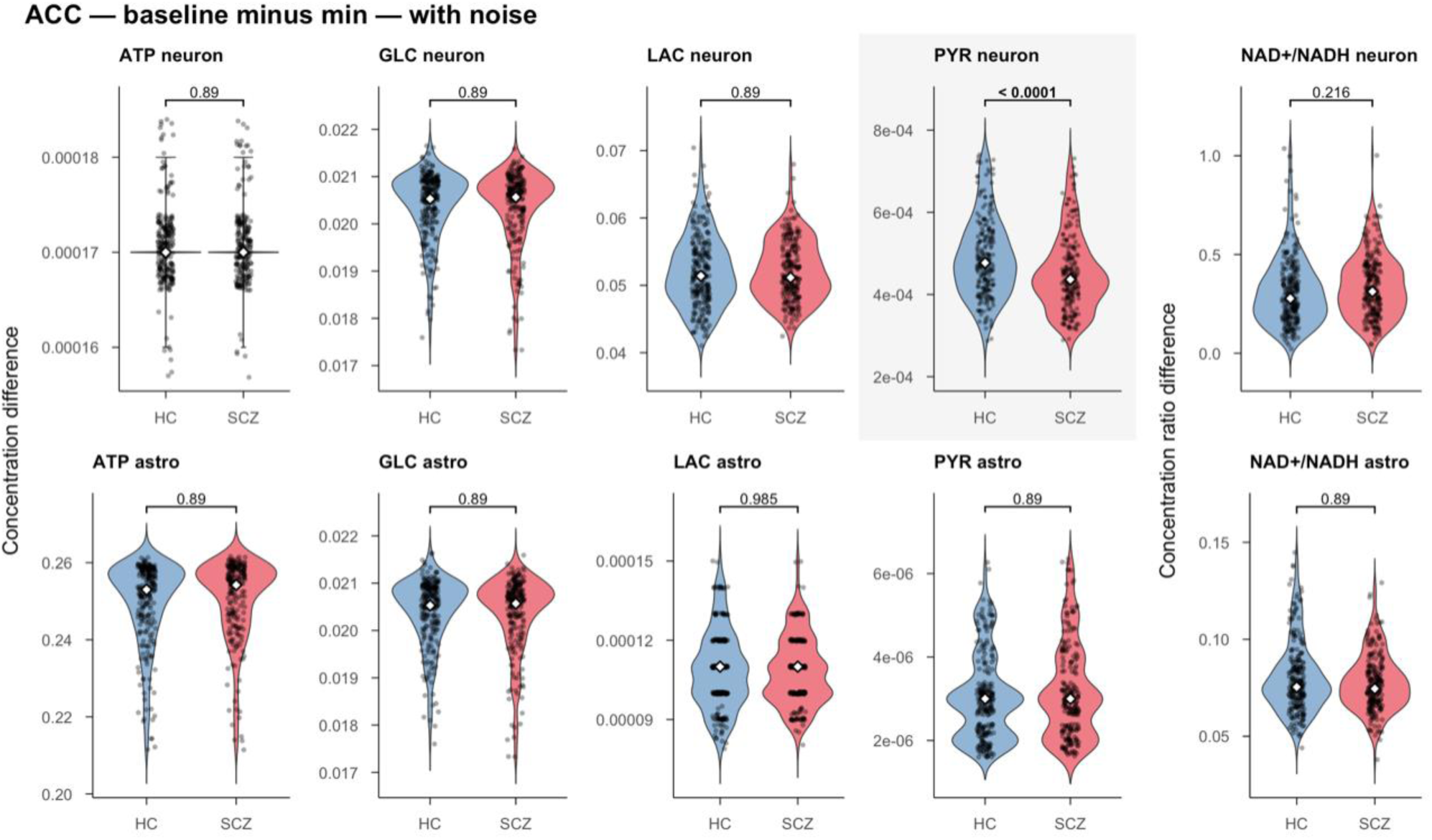
The difference between baseline and minimum concentration of metabolites in the ACC during the 1200 s time course with added noise. Outliers (n = 41) have been removed from the plots.

**Fig E.**
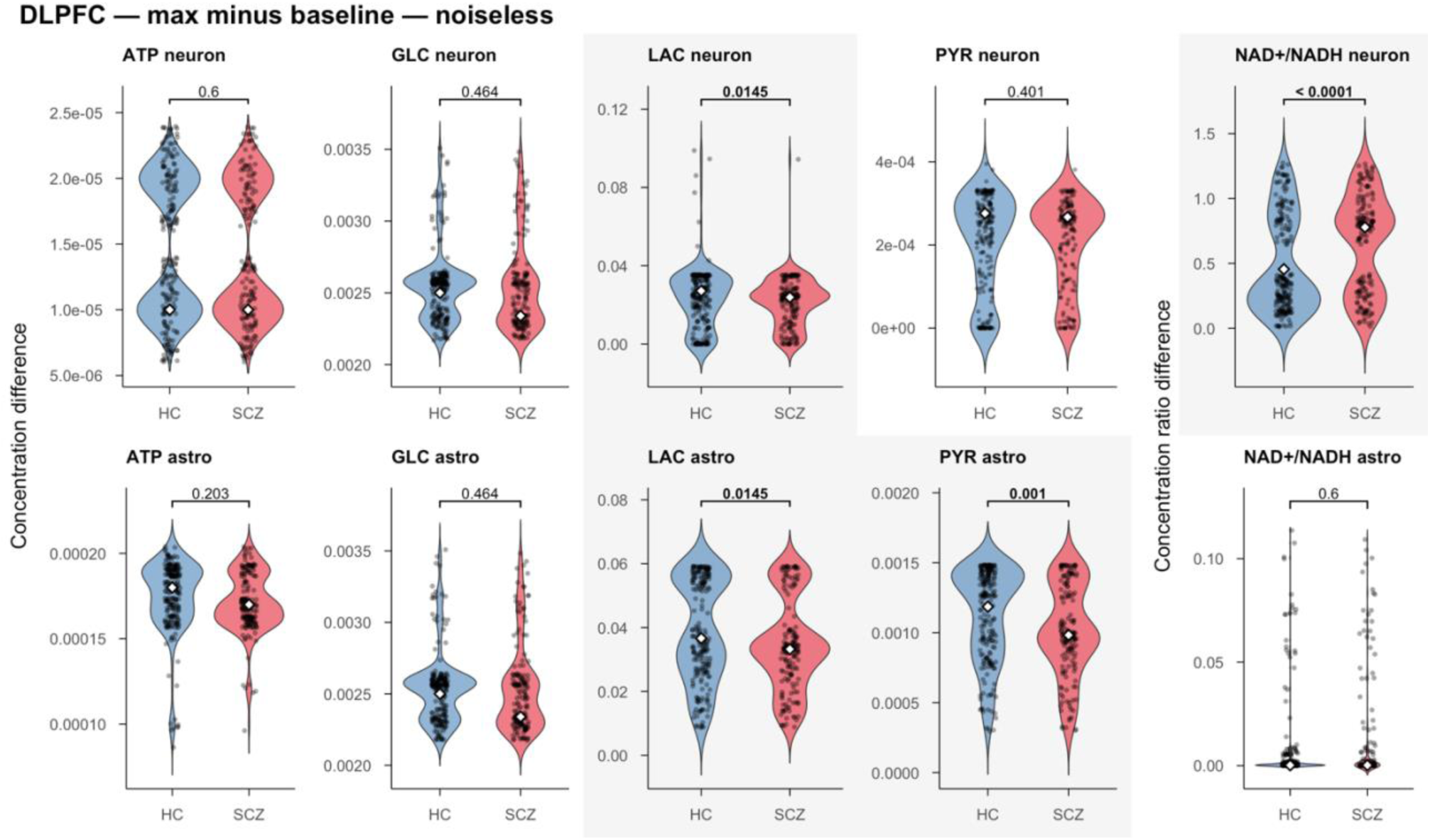
The difference between baseline and maximum concentration of metabolites in the DLPFC during the 1200 s time course using noiseless data. Outliers (n = 193) have been removed from the plots.

**Fig F.**
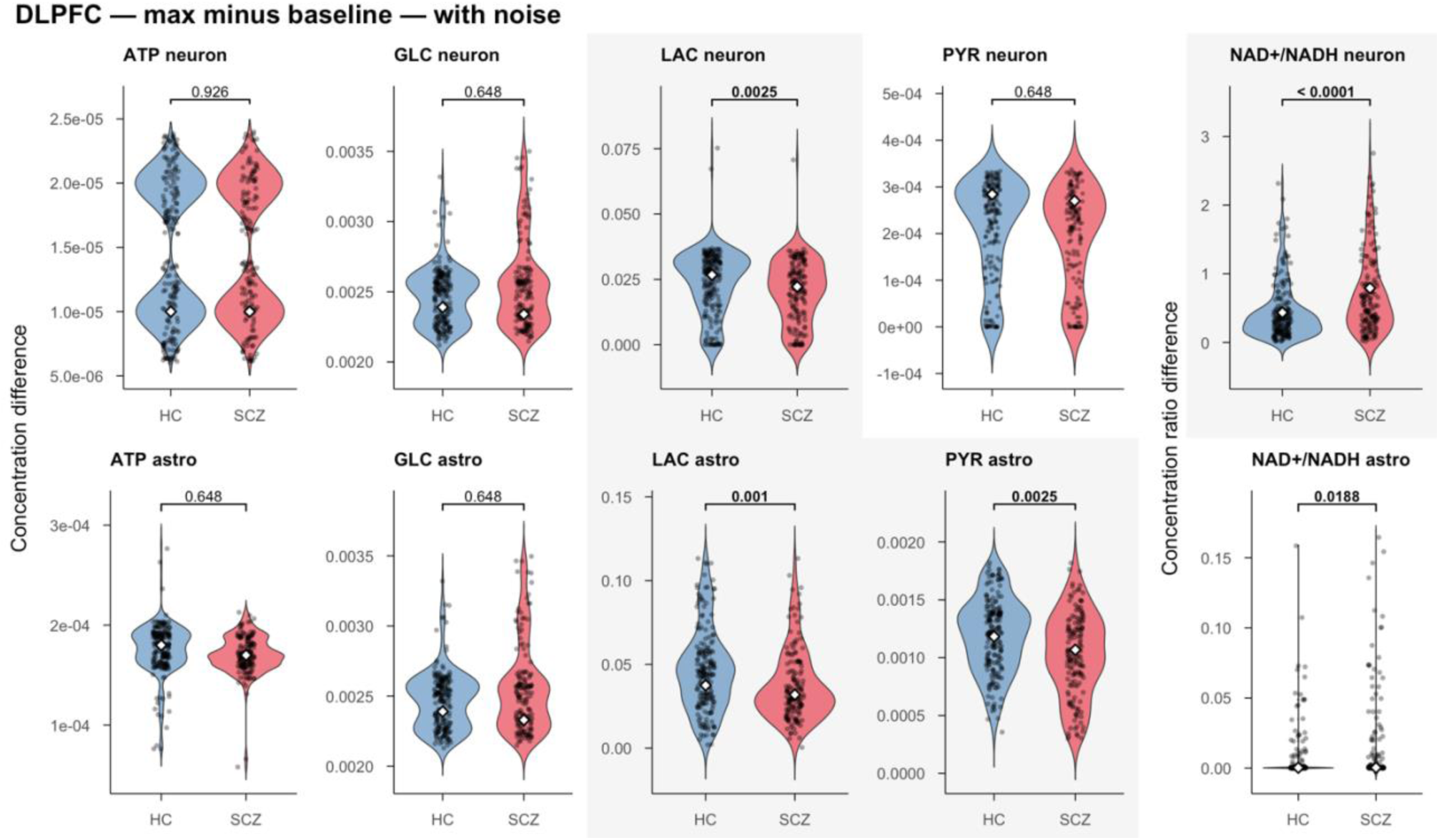
The difference between baseline and maximum concentration of metabolites in the DLPFC during the 1200 s time course with added noise. Outliers (n = 218) have been removed from the plots.

**Fig G.**
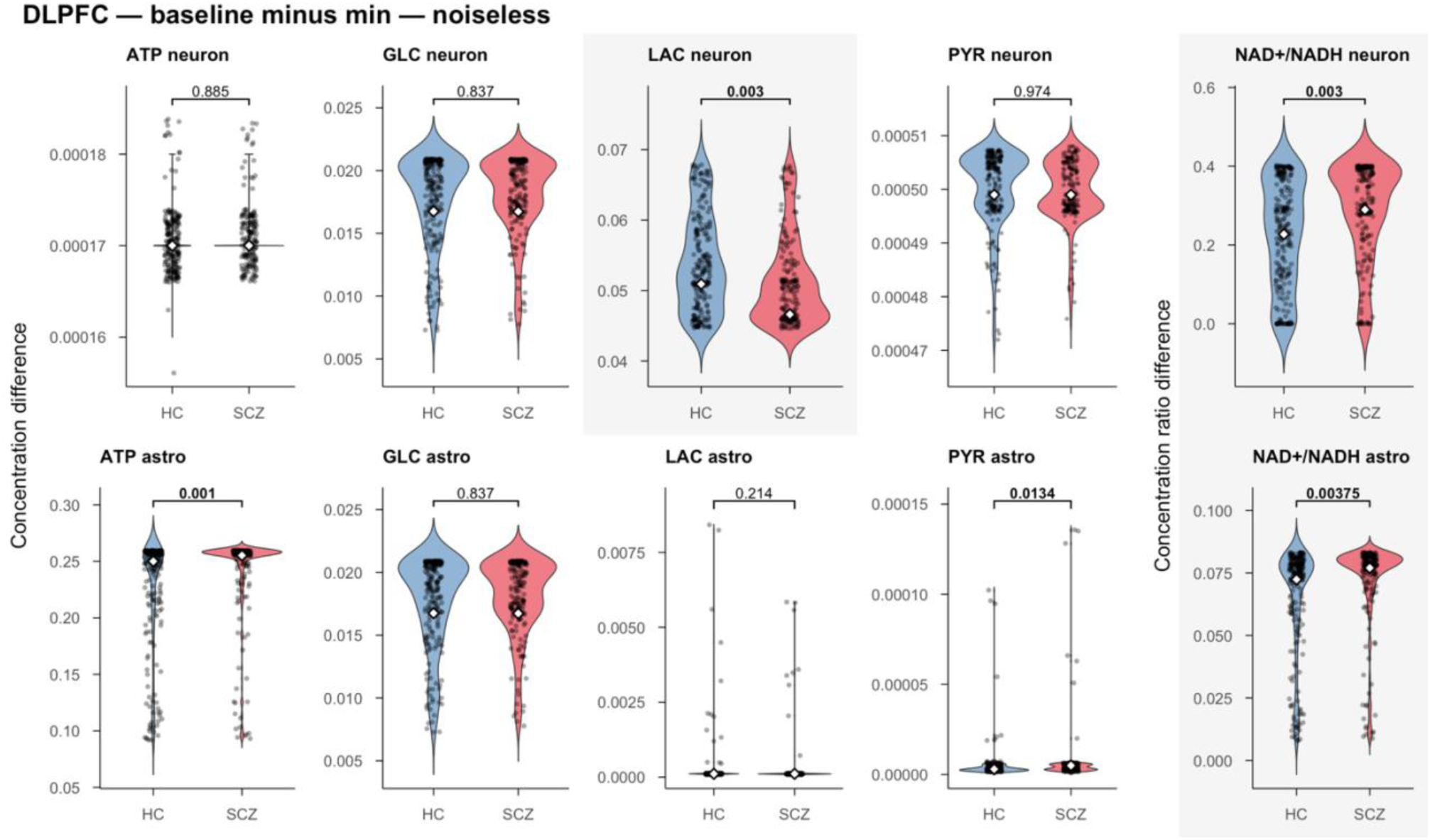
The difference between baseline and minimum concentration of metabolites in the DLPFC during the 1200 s time course using noiseless data. Outliers (n = 172) have been removed from the plots.

**Fig H.**
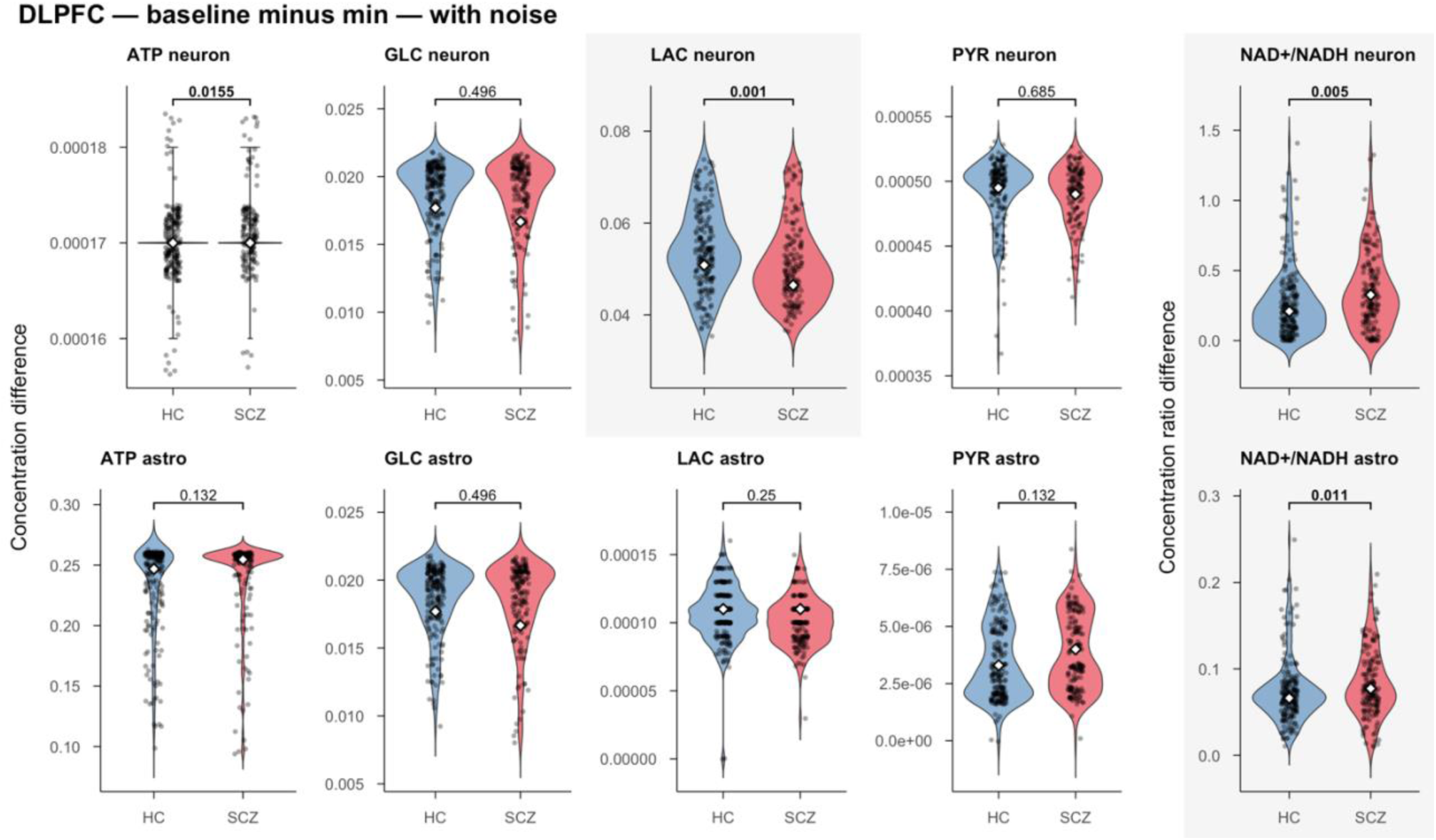
The difference between baseline and minimum concentration of metabolites in the DLPFC during the 1200 s time course with added noise. Outliers (n = 203) have been removed from the plots.

## REFERENCES

[1] McGrath J, Saha S, Chant D, Welham J. Schizophrenia: A Concise Overview of Incidence, Prevalence, and Mortality. Epidemiologic Reviews 2008;30:67–76.

[2] Jauhar S, Johnstone M, McKenna PJ. Schizophrenia. The Lancet 2022;399:473–86.

[3] McCutcheon RA, Reis Marques T, Howes OD. Schizophrenia-An Overview. JAMA Psychiatry 2020;77:201–10.

[4] Mueser KT, McGurk SR. Schizophrenia. Lancet 2004;363:2063–72.

[5] Sullivan PF, Kendler KS, Neale MC. Schizophrenia as a complex trait: evidence from a meta-analysis of twin studies. Arch Gen Psychiatry 2003;60:1187–92.

[6] Hilker R, Helenius D, Fagerlund B, Skytthe A, Christensen K, Werge TM, et al. Heritability of Schizophrenia and Schizophrenia Spectrum Based on the Nationwide Danish Twin Register. Biol Psychiatry 2018;83:492–8.

[7] Trubetskoy V, Pardiñas AF, Qi T, Panagiotaropoulou G, Awasthi S, Bigdeli TB, et al. Mapping genomic loci implicates genes and synaptic biology in schizophrenia. Nature 2022;604:502–8.

[8] Schizophrenia Working Group of the Psychiatric Genomics Consortium. Biological insights from 108 schizophrenia-associated genetic loci. Nature 2014;511:421–7.

[9] Owen MJ, Legge SE, Rees E, Walters JTR, O’Donovan MC. Genomic findings in schizophrenia and their implications. Mol Psychiatry 2023;28:3638–47.

[10] Smeland OB, Frei O, Dale AM, Andreassen OA. The polygenic architecture of schizophrenia - rethinking pathogenesis and nosology. Nat Rev Neurol 2020;16:366–79.

[11] Schijven D, Kofink D, Tragante V, Verkerke M, Pulit SL, Kahn RS, et al. Comprehensive pathway analyses of schizophrenia risk loci point to dysfunctional postsynaptic signaling. Schizophrenia Research 2018;199:195–202.

[12] Horváth S, Mirnics K. Schizophrenia as a disorder of molecular pathways. Biol Psychiatry 2015;77:22–8.

[13] Henkel ND, Wu X, O’Donovan SM, Devine EA, Jiron JM, Rowland LM, et al. Schizophrenia: a disorder of broken brain bioenergetics. Mol Psychiatry 2022;27:2393–404.

[14] Harris JJ, Jolivet R, Attwell D. Synaptic energy use and supply. Neuron 2012;75:762–77.

[15] Bélanger M, Allaman I, Magistretti PJ. Brain energy metabolism: focus on astrocyte-neuron metabolic cooperation. Cell Metab 2011;14:724–38.

[16] Bonvento G, Bolaños JP. Astrocyte-neuron metabolic cooperation shapes brain activity. Cell Metab 2021;33:1546–64.

[17] Pruett BS, Meador-Woodruff JH. Evidence for altered energy metabolism, increased lactate, and decreased pH in schizophrenia brain: A focused review and meta-analysis of human postmortem and magnetic resonance spectroscopy studies. Schizophr Res 2020;223:29–42.

[18] Rowland LM, Pradhan S, Korenic S, Wijtenburg SA, Hong LE, Edden RA, et al. Elevated brain lactate in schizophrenia: a 7 T magnetic resonance spectroscopy study. Transl Psychiatry 2016;6:e967–e967.

[19] Sullivan CR, O’Donovan SM, McCullumsmith RE, Ramsey A. Defects in Bioenergetic Coupling in Schizophrenia. Biol Psychiatry 2018;83:739–50.

[20] Dudzik P, Lustyk K, Pytka K. Beyond dopamine: Novel strategies for schizophrenia treatment. Medicinal Research Reviews 2024;44:2307–30.

[21] Mäki-Marttunen T, Kaufmann T, Elvsåshagen T, Devor A, Djurovic S, Westlye LT, et al. Biophysical Psychiatry—How Computational Neuroscience Can Help to Understand the Complex Mechanisms of Mental Disorders. Front Psychiatry 2019;10:534.

[22] Aubert A, Costalat R, Valabrègue R. Modelling of the coupling between brain electrical activity and metabolism. Acta Biotheor 2001;49:301–26.

[23] Aubert A, Costalat R. A model of the coupling between brain electrical activity, metabolism, and hemodynamics: application to the interpretation of functional neuroimaging. Neuroimage 2002;17:1162–81.

[24] Aubert A, Pellerin L, Magistretti PJ, Costalat R. A coherent neurobiological framework for functional neuroimaging provided by a model integrating compartmentalized energy metabolism. Proc Natl Acad Sci U S A 2007;104:4188–93.

[25] Aubert A, Costalat R. Interaction between astrocytes and neurons studied using a mathematical model of compartmentalized energy metabolism. J Cereb Blood Flow Metab 2005;25:1476–90.

[26] Cloutier M, Bolger FB, Lowry JP, Wellstead P. An integrative dynamic model of brain energy metabolism using in vivo neurochemical measurements. J Comput Neurosci 2009;27:391–414.

[27] Winter F, Bludszuweit-Philipp C, Wolkenhauer O. Mathematical analysis of the influence of brain metabolism on the BOLD signal in Alzheimer’s disease. J Cereb Blood Flow Metab 2018;38:304–16.

[28] Hoffman GE, Bendl J, Voloudakis G, Montgomery KS, Sloofman L, Wang Y-C, et al. CommonMind Consortium provides transcriptomic and epigenomic data for Schizophrenia and Bipolar Disorder. Sci Data 2019;6:180.

[29] Kanehisa M, Goto S. KEGG: kyoto encyclopedia of genes and genomes. Nucleic Acids Res 2000;28:27–30.

[30] The UniProt Consortium, Bateman A, Martin M-J, Orchard S, Magrane M, Adesina A, et al. UniProt: the Universal Protein Knowledgebase in 2025. Nucleic Acids Research 2025;53:D609–17.

31. [31] R Core Team (2021). R: A language and environment for statistical computing. R Foundation for Statistical Computing, Vienna, Austria. https://www.R-project.org/.

[32] Love MI, Huber W, Anders S. Moderated estimation of fold change and dispersion for RNA-seq data with DESeq2. Genome Biol 2014;15:550.

[33] Newman AM, Steen CB, Liu CL, Gentles AJ, Chaudhuri AA, Scherer F, et al. Determining cell type abundance and expression from bulk tissues with digital cytometry. Nat Biotechnol 2019;37:773–82.

[34] Zhang Y, Chen K, Sloan SA, Bennett ML, Scholze AR, O’Keeffe S, et al. An RNA-sequencing transcriptome and splicing database of glia, neurons, and vascular cells of the cerebral cortex. J Neurosci 2014;34:11929–47.

[35] Hoops S, Sahle S, Gauges R, Lee C, Pahle J, Simus N, et al. COPASI—a COmplex PAthway SImulator. Bioinformatics 2006;22:3067–74.

[36] Malik-Sheriff RS, Glont M, Nguyen TVN, Tiwari K, Roberts MG, Xavier A, et al. BioModels—15 years of sharing computational models in life science. Nucleic Acids Research 2019:gkz1055.

[37] Bittar PG, Charnay Y, Pellerin L, Bouras C, Magistretti PJ. Selective Distribution of Lactate Dehydrogenase Isoenzymes in Neurons and Astrocytes of Human Brain. J Cereb Blood Flow Metab 1996;16:1079–89.

[38] Dunaway GA, Kasten TP, Sebo T, Trapp R. Analysis of the phosphofructokinase subunits and isoenzymes in human tissues. Biochem J 1988;251:677–83.

[39] Lee J-H, Liu R, Li J, Zhang C, Wang Y, Cai Q, et al. Stabilization of phosphofructokinase 1 platelet isoform by AKT promotes tumorigenesis. Nat Commun 2017;8:949.

[40] Townsend L, Pillinger T, Selvaggi P, Veronese M, Turkheimer F, Howes O. Brain glucose metabolism in schizophrenia: a systematic review and meta-analysis of 18FDG-PET studies in schizophrenia. Psychol Med 2023;53:4880–97.

[41] Karlsson M, Zhang C, Méar L, Zhong W, Digre A, Katona B, et al. A single-cell type transcriptomics map of human tissues. Sci Adv 2021;7:eabh2169.

[42] Koepsell H. Glucose transporters in brain in health and disease. Pflugers Arch 2020;472:1299–343.

[43] Wilpert N-M, Krueger M, Opitz R, Sebinger D, Paisdzior S, Mages B, et al. Spatiotemporal Changes of Cerebral Monocarboxylate Transporter 8 Expression. Thyroid 2020;30:1366–83.

[44] Davalieva K, Maleva Kostovska I, Dwork AJ. Proteomics Research in Schizophrenia. Front Cell Neurosci 2016;10:18.

[45] Sahay S, Pulvender P, Rami Reddy MVSR, McCullumsmith RE, O’Donovan SM. Metabolic Insights into Neuropsychiatric Illnesses and Ketogenic Therapies: A Transcriptomic View. IJMS 2024;25:8266.

[46] Clark D, Dedova I, Cordwell S, Matsumoto I. Altered proteins of the anterior cingulate cortex white matter proteome in schizophrenia. Proteomics Clin Appl 2007;1:157–66.

[47] Clark D, Dedova I, Cordwell S, Matsumoto I. A proteome analysis of the anterior cingulate cortex gray matter in schizophrenia. Mol Psychiatry 2006;11:459–70, 423.

[48] Martins-de-Souza D, Gattaz WF, Schmitt A, Rewerts C, Maccarrone G, Dias-Neto E, et al. Prefrontal cortex shotgun proteome analysis reveals altered calcium homeostasis and immune system imbalance in schizophrenia. Eur Arch Psychiatry Clin Neurosci 2009;259:151–63.

[49] Sullivan CR, Koene RH, Hasselfeld K, O’Donovan SM, Ramsey A, McCullumsmith RE. Neuron-specific deficits of bioenergetic processes in the dorsolateral prefrontal cortex in schizophrenia. Mol Psychiatry 2019;24:1319–28.

[50] Mirnics K, Middleton FA, Marquez A, Lewis DA, Levitt P. Molecular characterization of schizophrenia viewed by microarray analysis of gene expression in prefrontal cortex. Neuron 2000;28:53–67.

[51] Hakak Y, Walker JR, Li C, Wong WH, Davis KL, Buxbaum JD, et al. Genome-wide expression analysis reveals dysregulation of myelination-related genes in chronic schizophrenia. Proc Natl Acad Sci U S A 2001;98:4746–51.

[52] Batiuk MY, Tyler T, Dragicevic K, Mei S, Rydbirk R, Petukhov V, et al. Upper cortical layer–driven network impairment in schizophrenia. Sci Adv 2022;8:eabn8367.

[53] Ruzicka WB, Mohammadi S, Fullard JF, Davila-Velderrain J, Subburaju S, Tso DR, et al. Single-cell multi-cohort dissection of the schizophrenia transcriptome. Science 2024;384:eadg5136.

[54] Frame AK, Robinson JW, Mahmoudzadeh NH, Tennessen JM, Simon AF, Cumming RC. Aging and memory are altered by genetically manipulating lactate dehydrogenase in the neurons or glia of flies. Aging (Albany NY) 2023;15:947–81.

[55] Liu S, Zhang L, Fan X, Wang G, Liu Q, Yang Y, et al. Lactate levels in the brain and blood of schizophrenia patients: A systematic review and meta-analysis. Schizophrenia Research 2024;264:29–38.

[56] Saudubray J-M, Garcia-Cazorla A. An overview of inborn errors of metabolism affecting the brain: from neurodevelopment to neurodegenerative disorders. Dialogues Clin Neurosci 2018;20:301–25.

[57] Bitanihirwe BKY, Woo T-UW. Oxidative stress in schizophrenia: an integrated approach. Neurosci Biobehav Rev 2011;35:878–93.

[58] Stein A, Zhu C, Du F, Öngür D. Magnetic Resonance Spectroscopy Studies of Brain Energy Metabolism in Schizophrenia: Progression from Prodrome to Chronic Psychosis. Curr Psychiatry Rep 2023;25:659–69.

[59] Mäki-Marttunen T, Blackwell KT, Akkouh I, Shadrin A, Valstad M, Elvsåshagen T, et al. Genetic mechanisms for impaired synaptic plasticity in schizophrenia revealed by computational modeling. Proc Natl Acad Sci USA 2024;121:e2312511121.

[60] Kim Y, Vadodaria KC, Lenkei Z, Kato T, Gage FH, Marchetto MC, et al. Mitochondria, Metabolism, and Redox Mechanisms in Psychiatric Disorders. Antioxid Redox Signal 2019;31:275–317.

[61] Fizíková I, Dragašek J, Račay P. Mitochondrial Dysfunction, Altered Mitochondrial Oxygen, and Energy Metabolism Associated with the Pathogenesis of Schizophrenia. Int J Mol Sci 2023;24:7991.

[62] Magistretti PJ, Allaman I. A cellular perspective on brain energy metabolism and functional imaging. Neuron 2015;86:883–901.

## References

[1] M. Karlsson, C. Zhang, L. Méar, W. Zhong, A. Digre, B. Katona, E. Sjöstedt, L. Butler, J. Odeberg, P. Dusart, F. Edfors, P. Oksvold, K. von Feilitzen, M. Zwahlen, M. Arif, O. Altay, X. Li, M. Ozcan, A. Mardinoglu, L. Fagerberg, J. Mulder, Y. Luo, F. Ponten, M. Uhlén, C. Lindskog, ‘A single-cell type transcriptomics map of human tissues’ Sci Adv, vol. 7, no. 31, pp. eabh2169, Jul. 2021.

[2] M. T. J. Lowe, E. H. Kim, R. L. M. Faull, D. L. Christie, and H. J. Waldvogel, ‘Dissociated expression of mitochondrial and cytosolic creatine kinases in the human brain: a new perspective on the role of creatine in brain energy metabolism’, J Cereb Blood Flow Metab, vol. 33, no. 8, pp. 1295–1306, Aug. 2013.

[3] T. Zheng, D. Kotol, R. Sjöberg, N. Mitsios, M. Uhlén, W. Zhong, F. Edfors, and J. Mulder ‘Characterization of reduced astrocyte creatine kinase levels in Alzheimer’s disease’, Glia, vol. 72, no. 9, pp. 1590–1603, Sep. 2024.

[4] R. H. Andres, A. D. Ducray, U. Schlattner, T. Wallimann, and H. R. Widmer, ‘Functions and effects of creatine in the central nervous system’, Brain Research Bulletin, vol. 76, no. 4, pp. 329–343, Jul. 2008.

[5] E. Wyatt, R. Wu, W. Rabeh, H.-W. Park, M. Ghanefar, and H. Ardehali, ‘Regulation and Cytoprotective Role of Hexokinase III’, PLoS ONE, vol. 5, no. 11, p. e13823, Nov. 2010.

[6] P. G. Bittar, Y. Charnay, L. Pellerin, C. Bouras, and P. J. Magistretti, ‘Selective Distribution of Lactate Dehydrogenase Isoenzymes in Neurons and Astrocytes of Human Brain’, J Cereb Blood Flow Metab, vol. 16, no. 6, pp. 1079–1089, Nov. 1996.

[7] G. A. Dunaway, T. P. Kasten, T. Sebo, and R. Trapp, ‘Analysis of the phosphofructokinase subunits and isoenzymes in human tissues’, Biochem J, vol. 251, no. 3, pp. 677–683, May 1988.

[8] J.-H. Lee, R. Liu, J. Li, C. Zhang, Y. Wang, Q. Cai, X. Qian, Y. Xia, Y. Zheng, Y. Piao, Q. Chen, J. F. de Groot, T. Jiang, and Z. Lu, ‘Stabilization of phosphofructokinase 1 platelet isoform by AKT promotes tumorigenesis’, Nat Commun, vol. 8, no. 1, p. 949, Oct. 2017.

[9] Y. Wei, M. Lu, M. Mei, H. Wang, Z. Han, M. Chen, H. Yao, N. Song, X. Ding, J. Ding, M. Xiao, and G. Hu, ‘Pyridoxine induces glutathione synthesis via PKM2-mediated Nrf2 transactivation and confers neuroprotection’, Nat Commun, vol. 11, no. 1, p. 941, Feb. 2020.

[10] H. Koepsell, ‘Glucose transporters in brain in health and disease’, Pflugers Arch, vol. 472, no. 9, pp. 1299–1343, Sep. 2020.

[11] N.-M. Wilpert, M. Krueger, R. Opitz, D. Sebinger, S. Paisdzior, B. Mages, A. Schulz, J. Spranger, E. K. Wirth, H. Stachelscheid, P. Mergenthaler, P. Vajkoczy, H. Krude, P. Kühnen, I. Bechmann, and H. Biebermann, ‘Spatiotemporal Changes of Cerebral Monocarboxylate Transporter 8 Expression’, Thyroid, vol. 30, no. 9, pp. 1366–1383, Sep. 2020.

[12] K. Banki, E. Colombo, F. Sia, D. Halladay, D. H. Mattson, A. H. Tatum, P. T. Massa, P. E. Phillips, and A. Perl, ‘Oligodendrocyte-specific expression and autoantigenicity of transaldolase in multiple sclerosis’, J Exp Med, vol. 180, no. 5, pp. 1649–1663, Nov. 1994.

